# Cancer cell type-specific derepression of transposable elements by inhibition of chromatin modifier enzymes

**DOI:** 10.1101/2024.01.15.575744

**Authors:** Divyesh Patel, Ville Tiusanen, Päivi Pihlajamaa, Biswajyoti Sahu

## Abstract

The combination of immunotherapy and epigenetic therapy is emerging as a promising approach for cancer therapy. Epigenetic therapy can induce derepression of transposable elements (TEs) that play a major role in activation of immune response against cancer cells. However, the molecular mechanism of TE regulation by distinct chromatin modifier enzymes (CME) and in the context of p53 is still elusive. Here, we used epigenetic drugs to inhibit distinct CMEs in p53 wild-type and p53-mutant colorectal and esophageal cancer cells. We show that distinct TEs subfamilies are derepressed by inhibition of different CMEs in a cell-type specific manner with loss of p53 resulting in stronger TE derepression. We show that KAP1, a known repressor of TEs, associates with stronger derepression of specific TE subfamilies such as LTR12C, indicating that KAP1 also has an activating role in TE regulation in cancer cells upon co-inhibition of DNMT and HDAC. Co-inhibition of DNMT and HDAC activates immune response by inducing inverted repeat Alu expression, reducing ADAR1-mediated Alu RNA editing and inducing cell type-specific TE-chimeric transcript expression. Collectively, our study demonstrates that inhibition of different CMEs results in derepression of distinct TEs in cell type-specific manner and by utilizing distinct mechanistic pathways, providing insights for epigenetic therapies that could selectively enhance anti-tumor immunity in distinct cancer types.

## Introduction

Transposable elements (TEs) and their remnants represent more than half of the human genome[1, 2]. TEs serve as reservoir of gene regulatory elements[3, 4] and can act as cancer-specific enhancers exploited by tissue-specific transcription factors[5] and thus TE expression and transposition can be pathogenic[4, 6]. In normal cells, TEs are epigenetically repressed by numerous mechanisms that include DNA methylation[4, 7], enrichment of repressive histone marks[4, 8], recruitment of Krüppel-associated box zinc finger (KRAB-ZNF)/KRAB-associated protein 1 (KAP1/TRIM28) complex[9, 10] and binding of p53[11]. KRAB-ZNFs bind to TEs and recruit KAP1 through KRAB domain[10, 12]. KAP1 in turn recruits the repressive chromatin modifier enzymes including heterochromatin protein HP1, histone H3K9 methyltransferase SETDB1, and the nucleosome remodeling and deacetylase (NuRD) complex[10, 13]. The role of KAP1 in repression of TEs is well characterized[9, 12]. However, recent studies have shown KAP1 recruitment to actively transcribed genes and its role in releasing paused Polymerase II[14–16]. However, the mechanism of KAP1 as transcriptional activator of TEs in cancer cells is not known.

The combination of epigenetic therapy and immune therapy have emerged as a major approach for cancer treatment to overcome the limitations of the immunotherapy, since epigenetic therapy can make cancer cells more responsive to the immunotherapy[17–19]. SETDB1 amplification in human tumors is associated with resistance to immune checkpoint blockade therapy (ICB) and loss of SETDB1 derepresses TEs with potential to encode viral proteins to activate immune response[20]. Epigenetic therapy such as inhibition of DNA methyltransferases (DNMTi) or co-inhibition of DNMT and histone deacetylases (DNMTi-HDACi) can activate cryptic promoters within TEs that can splice into nearby protein coding genes, resulting in TE-chimeric transcripts[21, 22]. TE-chimeric transcripts encode immunogenic antigens in cancer cells that can be targeted by immune therapy[21–23]. Previous studies have shown that human endogenous retrovirus (HERV) elements from the LTR12C subfamily derepressed by DNMTi or DNMTi-HDACi mainly contribute to immunogenic TE-chimeric transcripts[21, 22]. Moreover, inhibition of DNMT induces expression of TEs that can form double-stranded RNA (dsRNA) to activate type I/III interferon response [24–27]. Derepressed TE-derived dsRNA induces viral mimicry that can make cancer cells sensitive to immune therapy[25, 28]. However, the molecular mechanism of TE derepression by distinct epigenetic therapy and cell-type specific regulation is poorly understood.

In this study, we show that distinct TE subfamilies are derepressed by inhibition of different CMEs in a cell type-specific manner, and that co-inhibition of DNMT and HDAC has an synergistic effect on TE depression. We show that complete loss of p53 or mutant p53 in cancer cells result in stronger derepression of TEs after inhibition of CMEs. Importantly, specific TE subfamilies derepressed by co-inhibition of DNMT and HDAC are bound by KAP1, a known repressor of TEs, along with enrichment of active enhancer marks such as H3K27ac. SETDB1 inhibition (SETDB1i) induces expression of TE-derived chimeric transcript expression in a cell type-specific manner. Moreover, our results show that co-inhibition of DNMT and HDAC activates immune response by (i) inducing inverting repeat Alu expression with concomitant loss of ADAR1 mediated Alu RNA editing and (ii) inducing TE-chimeric transcript formation. In summary, we describe epigenetic mechanism for co-inhibition of DNMT-HDAC induced TE derepression which contributes to activation of immune response.

## Results

### Inhibition of CMEs results in derepression of common and cell type-specific TE subfamilies

To study epigenetic regulation of TEs by CMEs in cancer cell type-specific manner and in the context of p53, we utilized cancer cell lines representing two endodermal origin cancers, namely colon and esophageal cancers, both reported to have high rates of somatic retrotransposition[29]. Specifically, three cell lines with different p53 status were used: the GP5d colon adenocarcinoma cells (henceforth GP5d) that express wild-type p53, a variant of GP5d cells with p53-depletion (p53-KO)[30], and OE19 esophageal cancer cells harboring a mutant p53 conferring loss of oligomerization activity[31]. In the experiments, each cell line was treated with the following CME inhibitors (CMEi): Decitabine (DAC) for DNMT inhibition (DNMTi), SB939 for HDAC inhibition (HDACi), Mitramycin A for SETDB1 inhibition (SETDB1i), and a combination of DAC and SB939 for co-inhibition of DNMT and HDAC (DNMTi-HDACi) (**Fig. 1a**). Following the treatments, RNA-seq was used to measure the expression of TEs and ChIP-seq for KAP1 and active histone marks to delineate the epigenetic changes associated to the derepressed TEs (**Fig. 1a**). Analysis of RNA-seq data using SQuIRE[32] pipeline combined with TEtranscripts[33] and Telescope[34] enables comparison of TE expression at the subfamily and individual locus level, respectively (**Fig. 1a**).

**Fig. 1.**
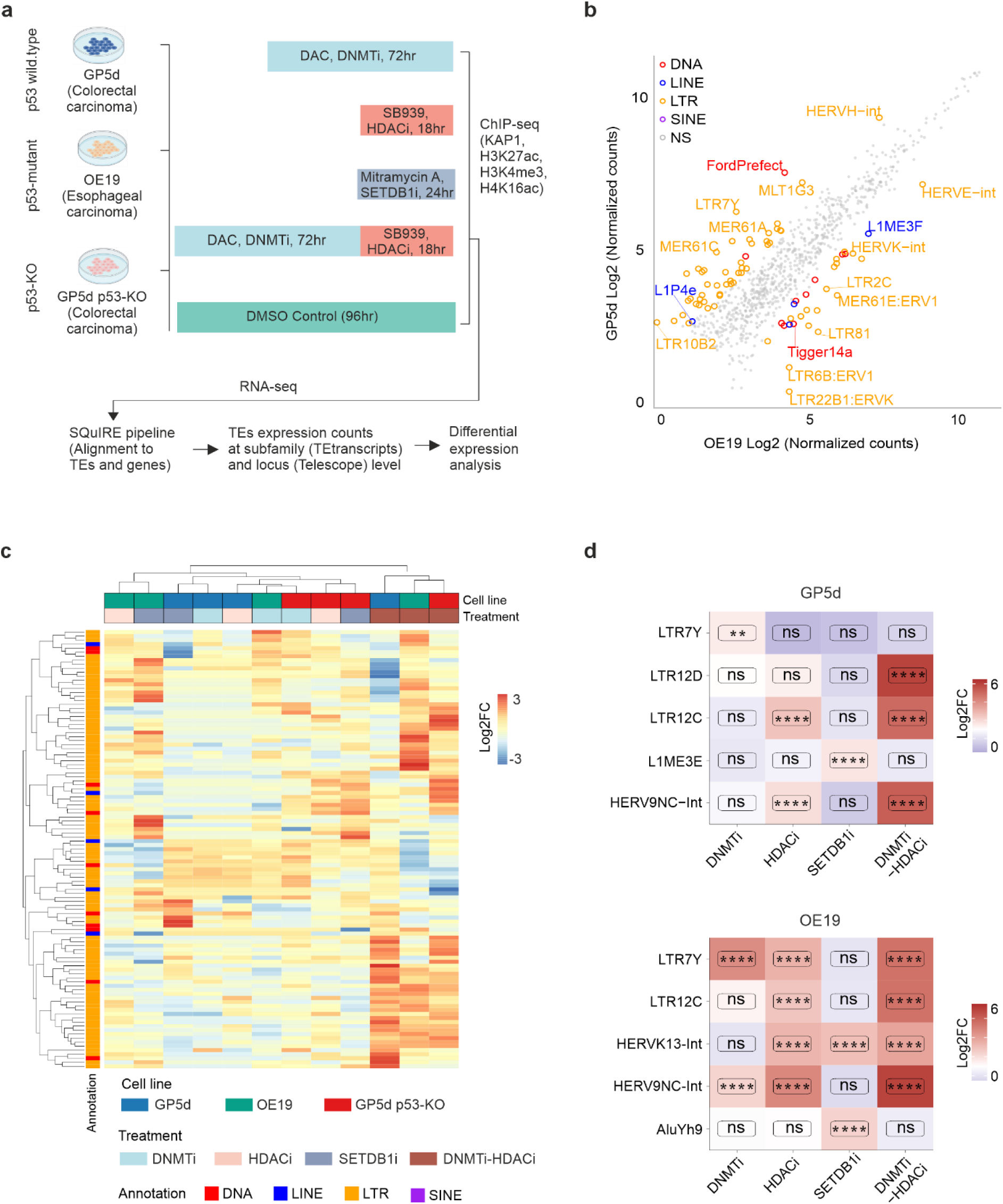
Common and cell type-specific TE subfamilies derepressed by inhibition of CMEs. a) Schematic representation of the CME inhibitor treatments and the analysis workflow pipeline. b) Differentially expressed TE subfamilies between DMSO treated GP5d and OE19 cells. Scatter plot shows normalized RNA-seq read counts for TE subfamilies. Differentially expressed TE subfamilies are labeled by TE class. c) Comparison of differentially expressed TE subfamilies induced by inhibition of CMEs in three cell lines. Differential expression analysis was performed by DESeq2. The heatmap shows Log2 fold change (FC) for TE subfamilies with absolute Log2FC > 2.5 (treatment vs vehicle-treated cells from the same cell line) and adjusted p-value < 0.05 in at least one CME treatment in at least one cell lines. Rows and columns are clustered with hierarchical clustering. d) Distinct TE subfamilies derepressed by inhibition of different CMEs in GP5d and OE19 cells. Expression changes for TE subfamilies (Log2FC) were compared between different CME treatments in GP5d (left panel) and OE19 (right panel) cells. Significance symbols: **** indicates p < 0.0001, ***p < 0.001, **p < 0.01, *p < 0.05, ns = non-significant |Log2FC|<1.5 or p > 0.05.

Comparative expression analysis between vehicle-treated GP5d and OE19 cells revealed cell type-specific expression of distinct TE subfamilies, most of the differentially expressed TEs belonging to the long terminal repeat (LTR) family (**Fig. 1b**). This is in agreement with previous reports from us and others showing differential TE expression between different cancer types [5, 35]. The observed differences partly reflect the different p53 status of these cell lines, since several LTRs known the be enriched for p53-binding sites such as LTR10b and MER61 elements[36] showed higher expression in GP5d cells compared to OE19 with mutated p53 (**Fig. 1b**). However, one of the MER61 subfamilies, MER61E, was upregulated in OE19 cells as well as LTRs from HERVK-int, HERVE-int and LTR22B1 subfamilies (**Fig. 1b**).

Next, we analyzed how the three cell lines representing colon and esophageal cancers respond to CME inhibition. Analysis of differentially expressed TE subfamilies induced by inhibition of distinct CMEs revealed that the overall effect of DNMTi on TE expression was weaker compared to HDACi and SETDB1i (**Supplementary Figure 1a**). Importantly, co-inhibition of DNMT and HDAC synergistically upregulated the expression of TE subfamilies compared to inhibition of DNMT or HDAC alone in all three cell lines (**Supplementary Figure 1a**), as has also been reported in other cancer types[21, 22, 37]. The p53 status had a clear effect on differential TE expression induced by CME inhibition, as higher number of TE subfamilies were differentially expressed in OE19 and GP5d p53-KO cells compared to GP5d cells (**Supplementary Figure 1a**). To analyze whether CMEi treatments induce similar or distinct changes in different cell types, we compared the expression of TE subfamilies upon CME inhibition between three cell lines (see Methods for details). A total of 109 TE subfamilies were differentially expressed by at least one CMEi treatment in at least one cell line, 94 of which were LTRs (**Fig. 1c, Supplementary Figure 1b, and Supplementary Data 1**). Hierarchical clustering of the differentially expressed TE subfamilies revealed that DNMTi-HDACi-treated samples from all three cell lines clustered together, whereas the treatments with each CME inhibitor alone resulted in more variable expression changes in different cell lines (**Fig. 1c**). Collectively, we observed two distinct patterns of TE derepression: (i) TE subfamilies that are similarly derepressed in all three cell lines and (ii) TE subfamilies with cell type-specific response to CME inhibition (**Fig. 1c; Supplementary Figure 1b**). Majority of LTR12 family members and HERV9 elements were derepressed in all three cell lines by DNMTi-HDACi treatment (**Supplementary Figure 1b**), but we also observed distinct subfamilies strongly derepressed in different cell types upon DNMTi-HDACi treatment, such as LTR10B1, LTR47A2, and HERVI-int in GP5d cells, and HERVK, MER41-int, MER41C, and HERVK13-int in OE19 cells (**Supplementary Figure 1b**). We also observed TE subfamilies derepressed by DNMTi, HDACi or SETDB1i treatments alone. Representative examples of the TE subfamilies derepressed by distinct CMEi in cell type-specific manner are shown in **Fig. 1d**. For example, LTR7Y elements were derepressed by DNMTi in GP5d cells, while DNMTi, HDACi and DNMTi-HDACi derepressed LTR7Y in OE19 (**Fig. 1d**). SETDB1i derepressed L1ME3E subfamily belonging to long interspersed nuclear elements (LINE) in GP5d cells and HERVK13-int and AluYh9 short interspersed nuclear elements (SINE) in OE19 cells (**Fig. 1d**). LTR12C and HERV9NC-int showed stronger depression by DNMTi-HDACi than HDACi alone both in GP5d and OE19 cells (**Fig. 1d**). Collectively, our results show that TE subfamilies are under distinct epigenetic regulation, and that inhibition of distinct CMEs results in derepression of TE subfamilies both in common and in a cell type-specific manner.

#### DNMT and HDAC co-inhibition synergistically derepresses individual TE loci

To study the effect of inhibition of different CMEs on TEs expression at individual locus level, we performed differential expression analysis for TE expression counts obtained using Telescope[34]. Similar to changes in expression observed at TE subfamily level, a greater number of TE loci showed differential expression pattern by distinct inhibitor treatments in OE19 cells compared to GP5d cells (**Fig. 2a and Supplementary Figure 1a**). Moreover, HDACi and SETDB1i resulted in stronger derepression of TE loci compared to DNMTi with at least 12 times greater number of differentially expressed TE loci in both GP5d and OE19 cells (**Fig. 2a**). Similarly, stronger derepressive effect was induced by co-inhibition of DNMT and HDAC when compared to either DNMTi or HDACi alone in both GP5d and OE19 cells. Comparison of major classes of TEs (DNA, LINE, LTR, and SINE) among the derepressed TE loci revealed that different TE classes were derepressed by inhibition of distinct CMEs in cell type-specific manner. In OE19 cells, LINEs and SINEs constitute the majority of depressed elements after all CMEi treatments representing 32-38% and 22-39% of the derepressed elements, respectively (**Fig. 2b, Supplementary Data 2**). In GP5d cells, by contrast, different CMEi treatments result in more variable response in TE derepression and a larger proportion of the derepressed elements belong to LTRs. In particular, DNMTi as well as DNMTi-HDACi induced LTR derepression (40 and 41% of the total elements, respectively), whereas only 19% of the SETDB1i-induced elements were LTRs and 42% SINEs (**Fig. 2b, Supplementary Data 2**). Overlap analysis of derepressed TE loci shows little overlap between different CMEi treatments in both GP5d and OE19 cells (**Fig. 2c**), suggesting that each chromatin modifier contributes to the control of distinct set of TE loci. The derepression of a greater number of TE loci by DNMTi-HDACi than DNMTi, HDACi and SETDB1i treatments alone in GP5d as well as OE19 cells (**Supplementary Data 2**). Collectively, our results demonstrate distinct patterns of TE regulation by different chromatin modifier enzymes, and that co-inhibition of DNMT and HDAC enzymes synergistically derepresses individual TE loci in a cancer cell-specific manner.

**Fig. 2.**
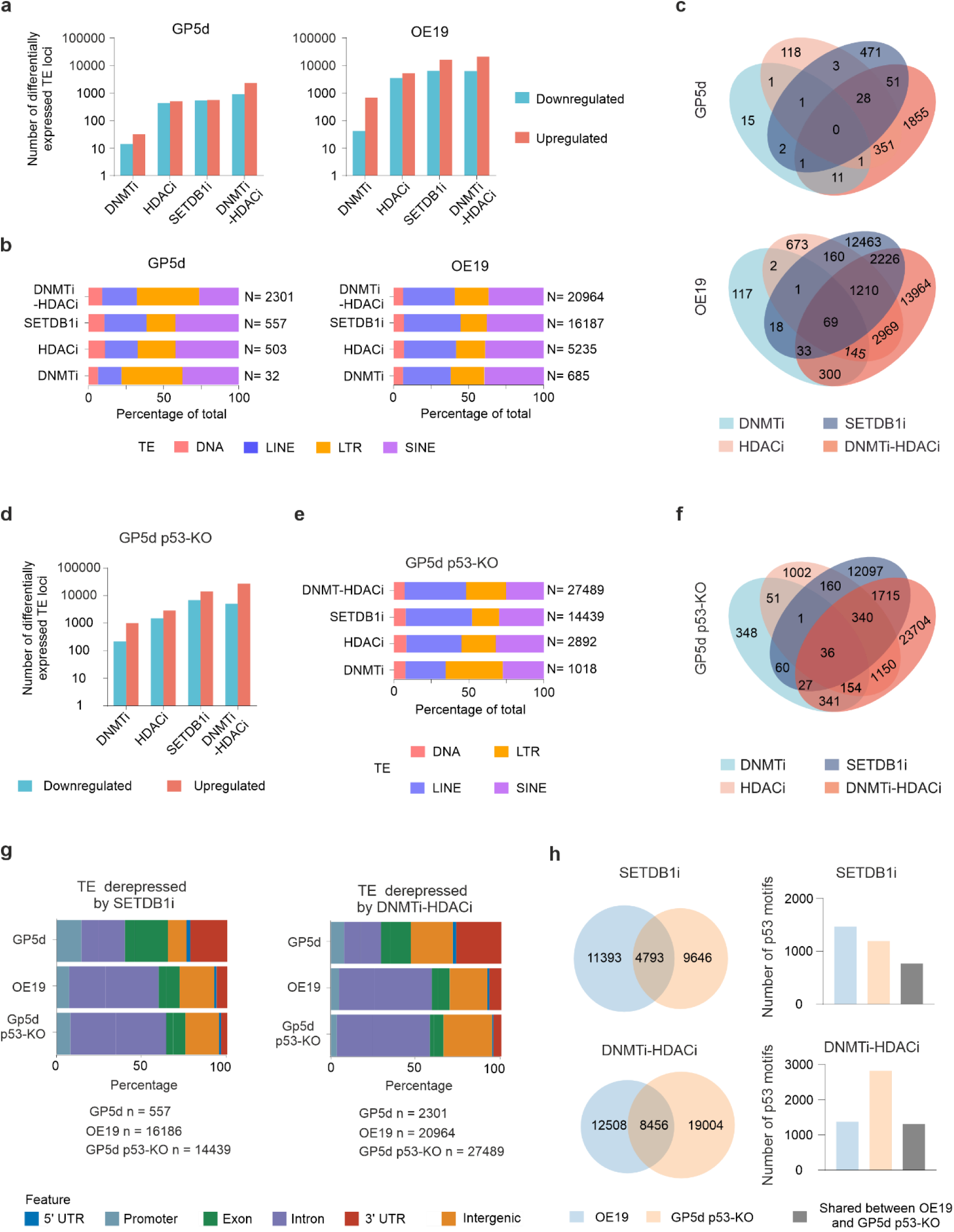
DNMT and HDAC co-inhibition synergistically derepresses TEs. a) Number of differentially expressed TE loci induced by inhibition of CMEs in GP5d and OE19 cells. TE loci that meet threshold criteria of Log2FC > 1.5 and adjusted p-value < 0.05 compared to vehicle-treated cells are considered as differentially expressed (DESeq2). b) The proportion of derepressed TE loci by inhibition of CMEs belonging to major TE classes in GP5d and OE19 cells. Derepressed TE loci were labeled by TE class and their counts presented as percentage of total. Numbers represent total derepressed TE loci. c) Overlap of derepressed individual TE loci by inhibition of CMEs in GP5d and OE19 cells. d) Number of differentially expressed TE loci by inhibition of distinct CMEs in GP5d p53-KO cells. Differentially expressed TEs were filtered out as described in Fig. 2a. e) The proportion of derepressed TE loci by inhibition of CMEs belonging to major TE classes in GP5d p53-KO cells. Derepressed TE loci were labeled by TE class and their counts presented as percentage of total. Numbers represent total derepressed TE loci. f) Overlap of derepressed individual TE loci by inhibition of CMEs in GP5d p53-KO cells. g) Genomic distribution of TEs derepressed by SETDB1i (left panel) and DNMTi-HDACi (right panel) in GP5d, OE19 and GP5d p53-KO cells. h) Overlap of TE loci derepressed by SETDB1i and DNMTi-HDACi in OE19 and GP5d p53-KO cells (left panel). TE derepressed by SETDB1i and DNMTi-HDACi were grouped into unique to OE19, unique to GP5d p53-KO and shared between OE19 and GP5d p53-KO. Enrichment of p53 motifs were compared between three groups (right panel).

### p53 loss results in stronger derepression of TEs upon DNMT and HDAC co-inhibition

Due to the known role of p53 in transcriptional repression of TEs[11], we set out to study the role of p53 in cell type-specific TE regulation in cells treated with distinct CMEs. For this, we performed the CME inhibitor treatments in p53-depleted (p53-KO) GP5d cells and compared the TE expression pattern with GP5d and OE19 cells harboring wild-type and mutant p53, respectively. Overall, the effect of CME inhibition on differential expression of TEs was stronger in p53-KO GP5d cells as compared to GP5d and OE19 cells (**Fig. 2a,d**). Interestingly, LINEs represented a higher percentage of the derepressed TE loci in GP5d p53-KO cells compared to GP5d cells by inhibition of distinct CMEs (**Fig. 2e, Supplementary Data 2**), in agreement with earlier reports showing higher LINE-1 expression in p53-mutant tumors[38]. Similar to GP5d and OE19 cells, derepressed TE loci elicited by different CMEi treatments in GP5d p53-KO cells showed minimal overlap (**Fig. 2f**) and co-inhibition of DNMT and HDAC synergistically derepressed the larger number of TEs compared to any of the treatments alone (**Fig. 2d and Supplementary Data 2**).

Next, we compared genomic location of the TEs derepressed by different CME inhibitor treatments in GP5d, OE19 and GP5d p53-KO cells. Interestingly, more than half of the depressed TEs induced by SETDB1i and DNMTi-HDACi treatments reside in intronic regions in both OE19 and GP5d-p53 KO cells (**Fig. 2g, Supplementary Data 3**). In contrast, UTRs and exons collectively represented more than half of the derepressed TE loci by distinct CMEs treatments in GP5d cells, respectively (**Fig. 2g, Supplementary Figure 2a, Supplementary Data 3**). As OE19 and GP5d p53-KO cells do not have wild-type p53 and derepressed TEs from both cell lines showed a similar genomic distribution pattern, we next compared the derepressed TEs in these two cell lines. Almost 30% and 40% of the TEs derepressed by SETDB1i and DNMTi-HDACi, respectively, are shared between OE19 and GP5d p53-KO cells (**Fig. 2h**, left panel). Furthermore, motif enrichment analysis revealed that most of the derepressed TEs (more than 80%) do not have p53-motif (**Fig. 2h**, right panel), indicating a p53-independent genomic control of these TEs.

To further analyze the role of p53 in controlling the TEs, we compared TE expression patterns between the two GP5d cell lines with and without functional p53. LTR subfamilies such as MER61 and LTR14 with higher proportion of genomic copies with p53 response elements (p53-RE) were upregulated in GP5d cells compared to p53-KO GP5d cells (**Supplementary Figure 3a**). Importantly, the absence of p53 potentiates the effect of DNMT and HDAC co-inhibition as 71 TE subfamilies were derepressed in p53-KO GP5d cells compared to 40 in GP5d cells (**Supplementary Figure 3b**, left panel). Similarly, 10-fold larger number of TE loci were derepressed in p53-KO GP5d cells (n = 27489) compared to GP5d cells (n = 2301) by DNMTi-HDACi (**Supplementary Figure 3b**, right panel).

Next, we measured the effect of DNMT and HDAC inhibition on the expression of p53 itself. Interestingly, DNMTi-HDACi treatment resulted in p53 downregulation at transcript and protein levels in both GP5d and OE19 cells, commensurate with reduced H3K27ac enrichment at the p53 promoter (**Supplementary Figure 3c,d**). We also observed smaller size of the p53 protein (approximately 40 kDa) in OE19 cells compared to GP5d cells (53 kDa), which validates earlier reported non-sense mutation in the tetramerization domain of p53[31] (**Supplementary Figure 3d**). ChIP-seq signal for H3K4me3 showed stronger reduction in GP5d cells compared to OE19 cells, which is in agreement with stronger downregulation of p53 expression in GP5d cells compared to OE19 cells (**Supplementary Figure 3c**). Furthermore, comparison of counts for differentially expressed TE loci in GP5d, OE19 and p53-KO GP5d cells showed an inverse correlation between functional activity of p53 and derepressed TEs loci counts (**Fig. 2a, 2d, and Supplementary Figure 3e**). To study the functional consequence of reduced p53 level elicited by DNMT and HDAC co-inhibition, we compared the expression of known p53 target genes regulated by LTRs harboring p53-REs[39], such as DHX37 and TMEM12 genes controlled by LTR10E and LTR10B1 elements, respectively. Both LTR10 elements are enriched for p53 binding and H3K27ac mark and the gene promoters for both H3K27ac and H3K4me3 marks in GP5d cells. Interestingly, loss of H3K27ac enrichment at p53-RE containing LTR10 elements and reduction of H3K4me3 levels at target gene promoters was observed upon DNMTi-HDACi with concomitant downregulation of DHX37 and TMEM12 genes (**Supplementary Figure 4a-b**). Collectively, DNMTi-HDACi treatment has a stronger effect on TE derepression in cells without functional p53 compared to p53-null cells, but it also results in downregulation of p53 and candidate p53-target genes and is associated with concordant loss of active enhancer marks from LTR elements having p53-REs.

### Different epigenetic mechanisms govern derepression of TEs induced by inhibition of DNMT and HDAC

Among the four different CMEi treatments, the strongest effect on TE derepression was elicited by combination of DNMTi-HDACi. Thus, we used DNMTi-HDACi treatment to characterize epigenetic changes at the derepressed TE loci. For this, we performed ChIP-seq in GP5d and OE19 cells with and without co-inhibition of DNMT and HDAC for well-known active histone marks H3K27ac and H3K4me3, as well as for H4K16ac, a histone mark recently reported at TE enhancers in human embryonic stem cells[40]. Moreover, we performed ChIP-seq for KAP1, since motif enrichment analysis performed for the genomic sequences of the depressed TE loci revealed enrichment of motifs for KRAB-ZFPs such as ZNF460, ZNF135 and ZBTB6 (**Fig. 3a**), and KAP1 is known to be recruited to TEs by KRAB-ZFPs to form KRAB-ZNF/KAP1 repressive complex[10, 12]. Recently, we reported derepression of TEs belonging to LTR12C subfamily by co-inhibition of DNMT and HDAC in GP5d cells[5], and in agreement with this, we also observed LTR12C derepression at both subfamily and individual locus level in OE19 cells (**Fig. 1d and 3b**). A total of 499 and 861 LTR12C loci were derepressed in GP5d and OE19 cells, respectively. A majority of these are shared between the two cell lines, as around 88% of derepressed LTR12C in GP5d cells overlap with the ones in OE19 cells (**Fig. 3c**). Enrichment analysis for TF binding motifs revealed motifs for forkhead family TFs, as well as GATA, ERF and NFY TFs enriched within these LTR12C sequences in both GP5d and OE19 cells (**Supplementary Figure 5a**). Enrichment of NFY motifs in derepressed LTR12C sequences agrees with earlier studies showing NFYA binding to be associated with transcription initiation from LTR12C[5, 21].

**Fig. 3.**
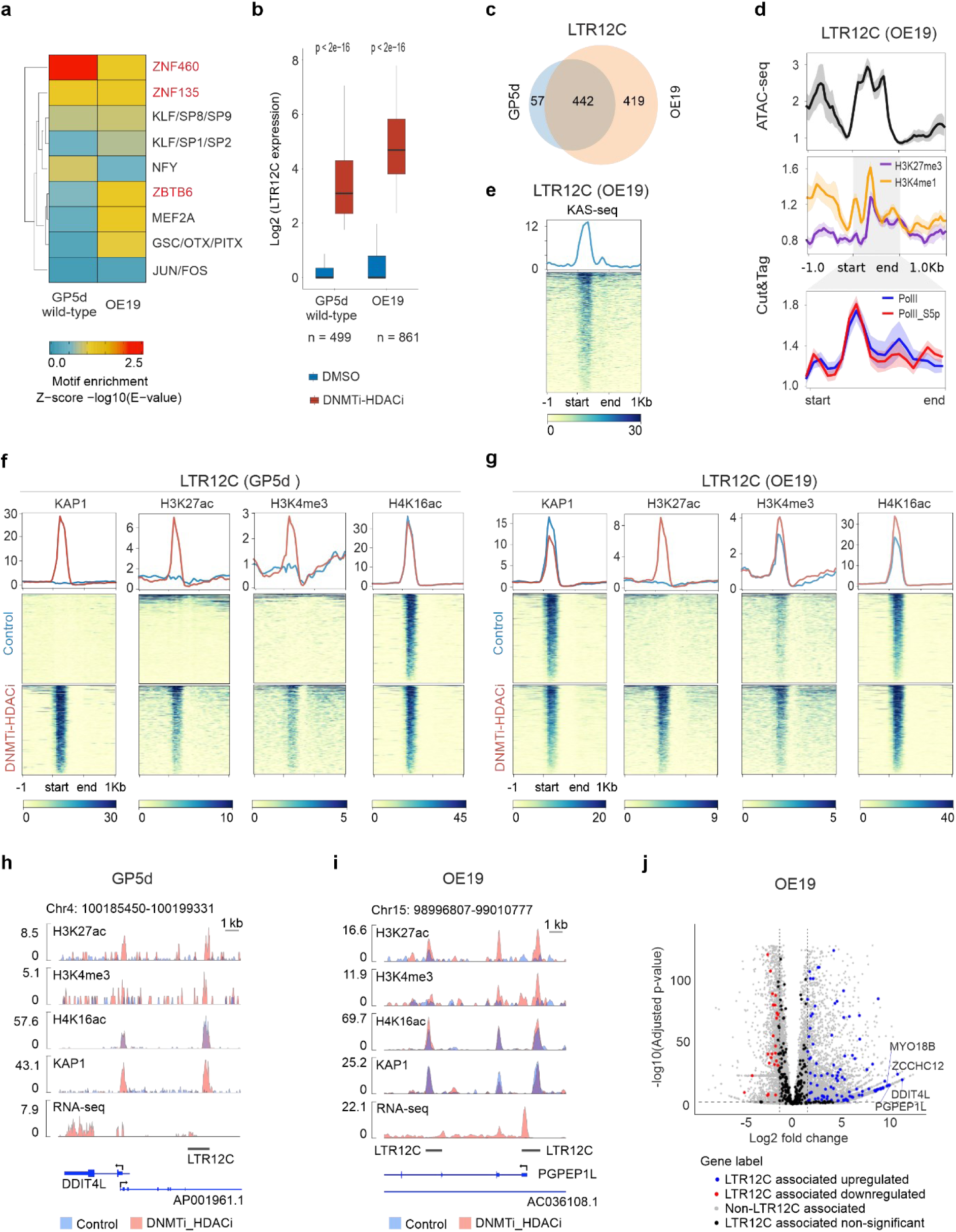
Different epigenetic mechanisms govern derepression of TEs by inhibition of DNMT and HDAC. a) TF motif enrichment at TE sequences derepressed by co-inhibition of DNMT and HDAC in GP5d and OE19 cells. After performing motif enrichment analysis for individual motifs, similar motifs were combined into motif clusters according to ref. [44]. The representative TF families are labeled on the right. b) Boxplots comparing the expression of derepressed LTR12C in GP5d and OE19 cells with and without co-inhibition of DNMT and HDAC (Wilcoxon paired test). c) Venn diagram showing overlap between LTR12C derepressed by co-inhibition of DNMT and HDAC in GP5d and OE19 (shown in Fig. 3b). d) Metaplots of ATAC-seq, CUT&TAG for H3K27me3 and H3K4me1, CUT&TAG for RNA-Pol II and Serine-5-phospohorylated RNA-Pol II in OE19 cells at LTR12C element derepressed by co-inhibition of DNMT and HDAC (shown in Fig. 3b). Derepressed LTR12C elements show a poised chromatin state in OE19 cells, whereas LTR12C in GP5d cells were enriched with repressive H3K27me3 marks (see **Supplementary Figure 5b**). e) Heatmap showing the KAS-seq signal in OE19 cells at LTR12C elements derepressed by co-inhibition of DNMT and HDAC (shown in Fig. 3B). f) Heatmaps showing the ChIP-seq signals for KAP1, H3K27ac, H3K4me3, and H4K16ac in GP5d cells with and without co-inhibition of DNMT and HDAC (shown in Fig. 3b). g) Heatmaps showing the ChIP-seq signals for KAP1, H3K27ac, H3K4me3, and H4K16ac in OE19 cells with and without inhibition of DNMT and HDAC (shown in Fig. 3b). h) Genome browser snapshot at a LTR12C element derepressed by co-inhibition of DNMT and HDAC in GP5d cells. Each panel show the ChIP-seq signals for H3K27ac, H3K4me3, H4K16ac, KAP1 and RNA-seq signal track for gene expression for both control and DNMTi-HDACi-treated cells. i) Genome browser snapshot at a LTR12C element derepressed by co-inhibition of DNMT and HDAC in OE19 cells. Each panel show the ChIP-seq signals for H3K27ac, H3K4me3, H4K16ac, KAP1 and RNA-seq signal track for gene expression for both control and DNMTi-HDACi treated cells. j) DNMTi-HDACi derepressed LTR12C elements in GP5d and OE19 cells. Volcano plots representing gene expression changes for derepressed LTR12C-associated genes for OE19 cells. Analysis of differentially expressed genes OE19 cells with DNMT and HDAC co-inhibition revealed significant up-regulation of genes in the vicinity of derepressed LTR12C elements (+/- 50kb).

Comparison of epigenetic profiles at derepressed LTR12C in GP5d and OE19 cells revealed that LTR12C elements are under different epigenetic regulation in GP5d and OE19 cells (**Fig. 3d and Supplementary Figure 5b**). In OE19 cells, derepressed LTR12C elements are enriched for ATAC-seq, as well as CUT&TAG signal for H3K27me3 and H3K4me1, suggesting a poised enhancer state[41] (**Fig. 3d**). Furthermore, CUT&TAG signal for RNA polymerase II and Ser-5-phosphorylation of RNA polymerase II showed pausing at LTR12C start sites (**Fig. 3d**). This is further supported by N3-kethoxal-assisted ssDNA sequencing (KAS-seq), a novel functional genomics assay to measure ssDNA, which represents transcriptionally engaged RNA Polymerase, transcribing enhancer, or non-canonical DNA structures[42]. In OE19 cells, derepressed LTR12C elements were enriched with single-stranded DNA signal from KAS-seq[43], which suggests transcriptionally engaged polymerase II at LTR12C (**Fig. 3e**). On the contrary, in GP5d cells derepressed LTR12C in GP5d cells were enriched for ATAC-seq and H3K27me3 mark but not for H3K4me1 signal (**Supplementary Figure 5b**). Collectively, these results suggest that LTR12C elements are under cell type-specific epigenetic regulation in open chromatin and are released from repression by DNMTi-HDACi treatment.

As most copies of LTR12C elements in the genome are truncated, it is difficult to distinguish the position of KAP1 binding between full-length LTR12C and truncated LTR12C elements. We filtered unique reads from KAP1 ChIP-seq (in GP5d DNMTi-HDACi and OE19 control samples) and OE19 KAS-seq that mapped to derepressed LTR12C elements and re-aligned the filtered reads to LTR12C consensus sequences. We observed KAP1 binding at the start sites of the LTR12C in both GP5d and OE19 cells (**Supplementary Figure 5c-d**). KAP1 binding also correlated with KAS-seq signal in OE19 cells (**Supplementary Figure 5d**). Next, we compared enrichment of ChIP-signal for KAP1 and active histone marks at derepressed LTR12C elements. In GP5d cells, KAP1 was recruited to the derepressed LTR12C together with gain of H3K27ac and H3K4me3 marks upon DNMTi-HDACi treatment (**Fig. 3f**). On the contrary, in OE19 cells KAP1 and H3K4me3 mark was already enriched at the LTR12C loci prior to inhibitor treatment (**Fig. 3g**), and co-inhibition of DNMT and HDAC only induced enrichment of H3K27ac marks (**Fig. 3g**). The enrichment of H4K16ac remained unaffected by co-inhibition of DNMT and HDAC in both cell lines (**Fig. 3f**). These results suggest cell type-specific differences in the epigenetic regulation and KAP1 function controlling the derepression of the TEs. Representative examples of derepressed LTR12C-mediated transactivation of nearby genes and cell type-specific patterns of KAP1 response to DNMTi-HDACi are shown for the DDIT4L gene in GP5d cells (**Fig. 3h**) and the PGPEP1L gene in OE19 cells (**Fig. 3i**). To support the role of LTR12C TEs as active enhancers controlling the expression of their nearby genes, we studied the effect of DNMTi-HDACi on the expression of genes located within (±50 kb) of the derepressed LTR12C elements in OE19 and GP5d cells. In total, 102 out of 132 and 70 out of 89 LTR12C-adjacent differentially expressed genes were upregulated by DNMTi-HDACi in OE19 and GP5d cells, respectively (**Fig. 3j and Supplementary Figure 5e**). These results suggest that derepression of LTR12C results in transactivation of nearby genes.

### Monoallelic KAP1 loss results in weaker derepression of LTR12C in GP5d cells

After observing KAP1 recruitment to derepressed LTR12C elements in GP5d cells by co-inhibition of DNMT and HDAC, we set out to delineate its role in regulation of TEs in DNMTi-HDACi-treated cells. For this, we used CRISPR-Cas9 system to delete one KAP1 allele from GP5d cells (GP5d KAP1^+/-^) and compared TE expression between DMSO- and DNMTi-HDACi-treated cells. Loss of single KAP1 allele reduced KAP1 expression at RNA and protein levels (**Fig. 4a and Supplementary Figure 5f**), although the reduction at RNA-level was stronger than protein level. As expected from known TE repressive role of KAP1, TE expression analysis revealed that co-inhibition of DNMT and HDAC resulted in stronger effect on TE derepression in GP5d KAP1^+/-^ cells compared to parental GP5d at subfamily level (**Fig. 4b**, left panel) as well as locus level (**Fig. 4b**, right panel). Interestingly, we also observed a small increase in the number of repressed TE subfamilies and TE loci by DNMTi-HDACi treatment in GP5d KAP1^+/-^ cells compared to GP5d cells (**Fig. 4b**, right panel), suggesting that KAP1 plays an activator role for these TEs in DNMTi-HDACi treated GP5d cells. The majority (83%) of the TEs derepressed by DNMTi-HDACi treatment in GP5d cells were similarly derepressed in GP5d KAP1^+/-^ cells (**Fig. 4c**). Comparison of major TE classes among the derepressed TEs showed that interspersed repeats, LINEs and SINEs, represented 5% larger proportion and LTRs 10% smaller proportion in GP5d KAP1^+/-^ cells compared to parental GP5d cells (**Fig. 4d**). Motif enrichment analysis for derepressed TEs showed motifs for KRAB-ZFPs such as ZNF460 and ZNF135 that were more enriched in TEs derepressed in GP5d cells, while motifs for RHOXF1, MAFK and ZNF257 were more enriched in TEs derepressed in GP5d KAP1^+/-^ (**Fig. 4e**). These differences in the enriched TF motifs suggest that different TFs might be associated to the derepressed TEs in GP5d cells depending on the KAP1 level.

**Fig. 4.**
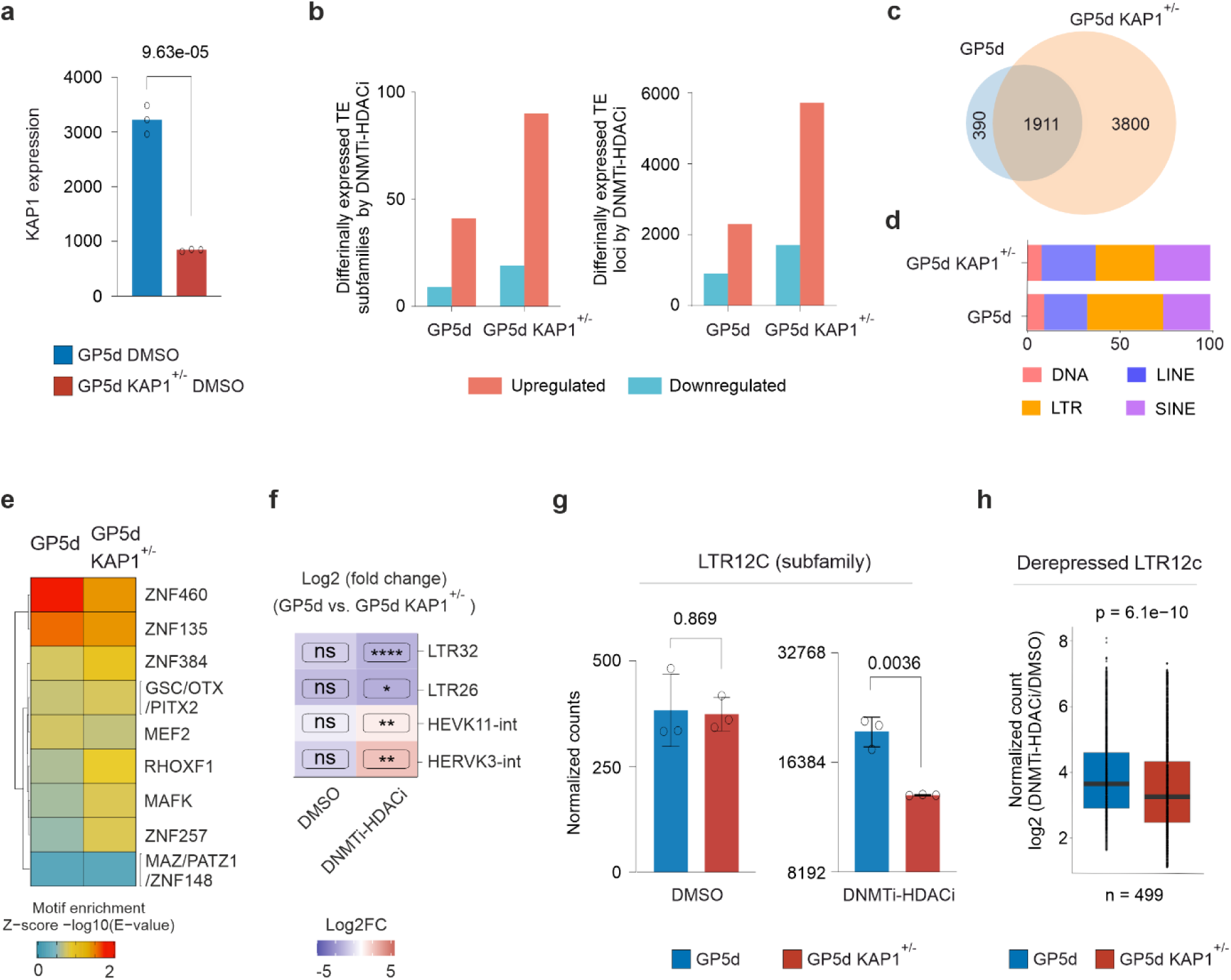
Monoallelic KAP1 loss results in weaker derepression of LTR12C in GP5d cells. a) Comparison of KAP1 gene expression in DMSO treated GP5d and GP5d KAP1^+/-^ cells (mean ± SD, individual data points for three biological replicates shown as dots; two-sided unpaired t-test). b) Number of differentially expressed TE subfamilies (left panel) and individual TE loci (right panel) by DNMTi-HDACi treatment in GP5d and GP5d KAP1^+/-^ cells. c) Venn diagram showing overlap between derepressed TE loci by DNMTi-HDACi treatment in GP5d and GP5d KAP1^+/-^ cells (from Fig. 4b, right panel). d) The proportion of derepressed TE loci by DNMTi-HDACi treatment belonging to major TE classes in GP5d and GP5d KAP1^+/-^ cells (from Fig. 4b, right panel). Derepressed TE loci are labeled by TE class and their counts are presented as percentage of total. Numbers represent total derepressed TE loci. e) Motif enrichment analysis for derepressed TEs by DNMTi-HDACi treatment in GP5d and GP5d KAP1^+/-^ cells (from Fig. 4b, right panel). Motif enrichment analysis was performed as described in Fig. 3a. f) Distinct TE subfamilies derepressed and repressed by DNMTi-HDACi treatment in GP5d and GP5d KAP1^+/-^ cells. Expression changes for TE subfamilies (Log2FC) were compared between GP5d and GP5d KAP1^+/-^ cells treated with DMSO or DNMTi-HDACi. (Significance symbols: **** indicates p < 0.0001, ***p < 0.001, **p < 0.01, *p < 0.05, ns = non-significant |Log2FC|<1.5 or p > 0.05). g) Comparison of LTR12C subfamily expression in GP5d and GP5d KAP1^+/-^ cells after DMSO or DNMTi-HDACi treatment. The normalized RNA-seq read counts for the LTR12C subfamily in DMSO treated cells (left panel) and DNMTi-HDACi treated cells (right panel) (mean ± SD, individual data points for three biological replicates shown as dots; two-sided unpaired t test). h) Comparison of derepressed LTR12C expression in GP5d and GP5d KAP1^+/-^ cells in DMSO or DNMTi-HDACi treated cells. The normalized LTR12C loci expression counts were compared between DNMTi-HDACi and DMSO treated (Wilcoxon paired test).

To characterize the repressive and activating roles of KAP1 in TE regulation in DNMTi-HDACi-treated cells, we compared the expression of TE subfamilies in GP5d and GP5d KAP1^+/-^ cells with and without co-inhibition of DNMTi and HDACi. LTR32 and LTR26 were repressed in GP5d KAP1^+/-^ cells compared to GP5d cells by DNMTi-HDACi treatment, whereas HERVK11-int and HERVK3-int showed higher derepression in GP5d KAP1^+/-^ cells compared to GP5d cells by DNMTi-HDACi treatment (**Fig. 4f**). Interestingly, loss of single allele of KAP1 also resulted in weaker derepression of LTR12C upon co-inhibition of DNMT and HDAC in GP5d KAP1^+/-^ cells compared to wild-type GP5d cells (**Fig. 4g**, right panel), while there was no difference in the LTR12C expression in the DMSO-treated cells. As expected from subfamily level expression, 77% (386 out of 499) of derepressed LTR12C (shown in **Fig. 3b**) showed weaker derepression in GP5d KAP1^+/-^ cells when compared to GP5d cells by DNMTi-HDACi treatment (**Fig. 4h**). Collectively, we observed a dual role for KAP1 in TE regulation in GP5d cells, that of transcriptional repressor as well as activator at distinct TE subfamilies upon co-inhibition of DNMT and HDAC.

### Inverted repeat Alu SINEs are derepressed by co-inhibition of DNMT and HDAC in GP5d and OE19 cells

Out of all derepressed TE loci in OE19, GP5d, and GP5d p53-KO cells, SINEs represented 37%, 26% and 25%, respectively, and most of the derepressed SINEs were Alu elements (**Fig. 2b and 2e**). Alu SINEs, specifically inverted-repeat Alu (IR-Alu) elements, are a major source of double-stranded RNA that can activate innate and adaptive immune response[24, 25]. To characterize transcription and epigenetic changes at IR-Alu elements by co-inhibition of DNMT and HDAC, we first selected the transcriptionally active IR-Alu from all IR-Alu elements present in the human genome (with sum of RNA-seq reads from DMSO and DNMT-HDACi samples ≥5, see Method for details, resulting in total of 4966, 83952, and 84606 active IR-Alus in GP5d, OE19, and GP5d p53-KO cells, respectively) and analyzed the changes in their expression between the DNMTi-HDACi- and vehicle-treated cells. Majority of the transcriptionally active IR-Alu showed increased expression by co-inhibition of DNMT and HDAC in all three cell lines (**Fig. 5a**), but the derepression was considerably stronger in OE19 and GP5d p53-KO cells (at least 20-fold higher) compared to GP5d cells. In OE19 and GP5d p53-KO cells, more than 70% of the derepressed IR-Alus reside in the intronic regions, which is in sharp contrast to GP5d cells in which around 50% of the depressed IR-Alus are located in 3’-UTRs (**Fig. 5b**). Next, we compared ChIP-seq signal for KAP1 and histone marks at derepressed IR-Alu elements. Increased KAP1 binding and H3K27ac ChIP-signal was observed at the derepressed IR-Alus in GP5d and OE19 cells (**Fig. 5c**). OE19 cells showed stronger gain for KAP1 occupancy and H3K27ac ChIP-signal compared to GP5d cells, consistent with stronger increase in IR-Alu expression in OE19 compared to GP5d cells (**Fig. 5c and 5a**). As derepressed IR-Alu have shown poor enrichment in gene promoter regions, we did not observe gain of promoter-associated H3K4me3 marks by DNMTi-HDACi treatment (**Supplementary Figure 6a).** We did not observe gain of ChIP-seq signal for H4K16ac at derepressed IR-Alus (**Supplementary Figure 6a**). These results demonstrate TE class- and cell type-specific epigenetic mechanisms controlling TEs in human cells. While H3K37ac enrichment consistently increased and H4K16ac levels remained constant upon DNMTi-HDACi treatment, KAP1 binding and H3K4me3 enrichment had more distinct patterns. In GP5d cells, KAP1 was recruited to both LTR12C and IR-Alu elements upon DNMTi-HDACi treatment, whereas in OE19 cells KAP1 was already bound to LTR12C elements in the control cells and only recruited to IR-Alus in the DNMTi-HDACi-treated cells. Active H3K4me3 mark was increased in the LTR12C elements but not in IR-Alus in DNMTi-HDACi-treated GP5d cells, and the levels remained constant for both LTR12C and IR-Alus in OE19 cells.

**Fig. 5.**
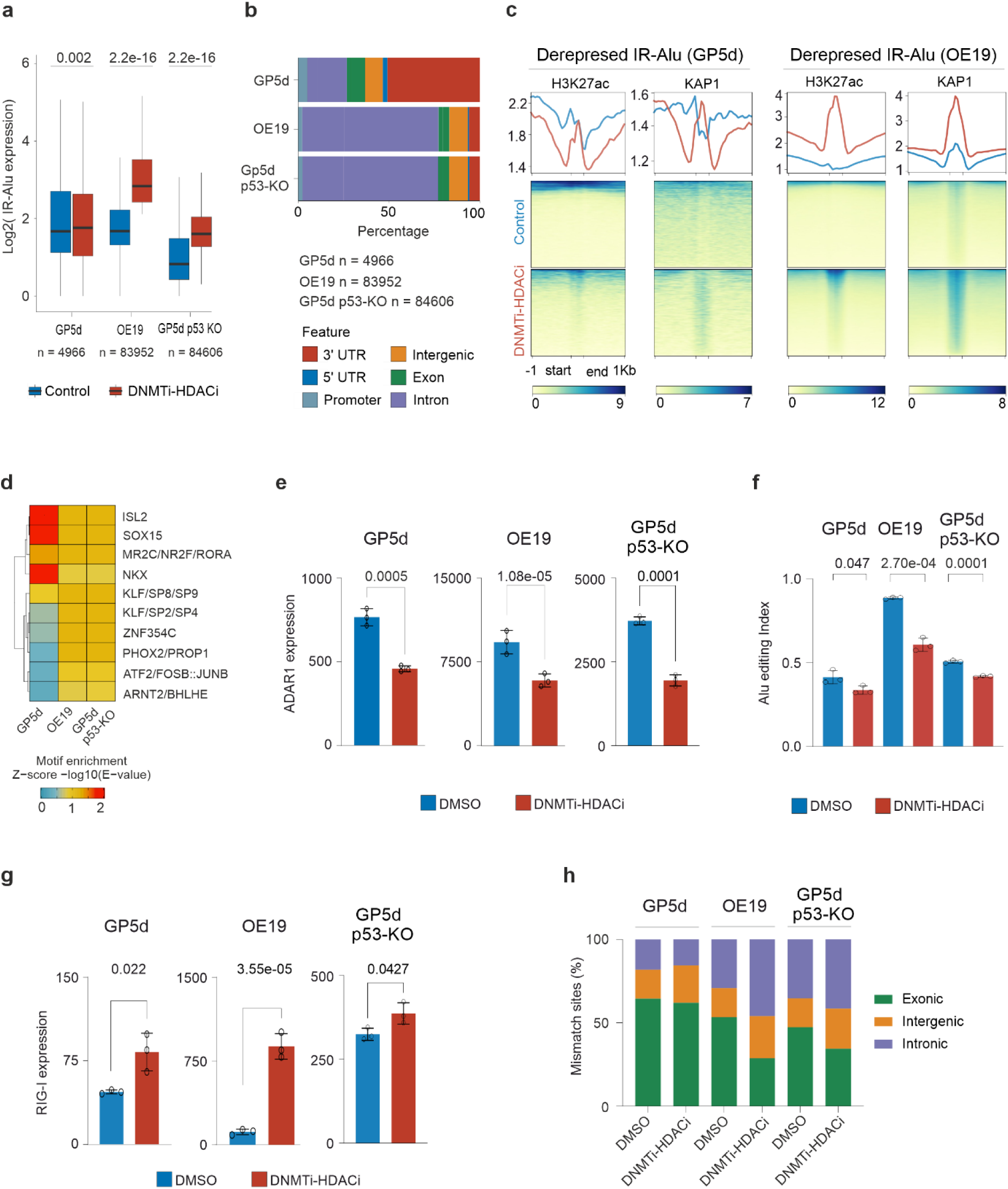
Inverted repeat Alu SINEs are derepressed by co-inhibition of DNMT and HDAC in GP5d and OE19 cells. a) Boxplots showing comparison of expression of transcriptionally active IR-Alu SINEs (Total sum of RNA-seq reads for DMSO and DNMT-HDACi ≥5, see methods for details) in GP5d, OE19 and GP5d p53-KO cells with and without co-inhibition of DNMT and HDAC. Majority of IR-Alu SINEs derepressed by co-inhibition of DNMT and HDAC (Wilcoxon paired test). b) Distribution of IR-Alu SINEs shown in Fig. 4a in annotated genomic regions. c) Heatmap showing the ChIP-seq signal for H3K27ac and KAP1 at IR-Alu SINEs in GP5d and OE19 cells with and without co-inhibition of DNMT and HDAC. d) Motif enrichment analysis for IR-Alu SINEs shown in Fig. 4a. Motif enrichment analysis was performed as described in Fig. 3a. e) Comparison of normalized RNA-seq read counts for ADAR1 gene in GP5d, OE19 and GP5d p53-KO cells with and without co-inhibition of DNMT and HDAC. f) Alu editing index (AEI) was calculated by using RNAeditingIndexer[47] tool on RNA-seq data. Bar plots showing AEI for GP5d, OE19 and GP5d p53-KO cells with and without co-inhibition of DNMT and HDAC. g) Bar plots compare RIG1 gene expression in GP5d, OE19 and GP5d p53-KO cells with and without co-inhibition of DNMT and HDAC. h) Alu RNA editing (A to G RNA editing) events were calculated for exons, intron and intergenic regions by using RNAeditingIndexer[47]. Distribution of Alu editing sites in GP5d, OE19 and GP5d p53-KO cells with and without co-inhibition of DNMT and HDAC.

Motif enrichment analysis of the sequences corresponding to the derepressed IR-Alu elements in GP5d, OE19 and GP5d p53-KO cells showed enrichment of NR2, RORA and SP5 motifs in all cell lines, but distinct TF motif enrichment patterns were observed as well. In GP5d cells, derepressed IR-Alus were enriched for ISL2, SOX15 and NKX motifs, whereas derepressed IR-Alus in OE19 and GP5d p53-KO cells were enriched for KLF/SP, ZNF354C, FOSB:JUNB and PHOX2/PROP1 motifs, suggesting cell type-specific *cis*-regulatory logic at these repeat elements at distinct genomic locations (**Fig. 5d**).

It has been previously reported that IR-Alu elements can form double stranded RNA (dsRNA) structures and that adenosine deaminases acting on RNA (ADAR1) enzyme[45] can act on dsRNA to alter gene regulation by post-transcriptional conversion of adenosine to inosine (A-to-I editing)[46]. ADAR1 enzymes edit dsRNA from IR-Alus to inhibit activation immunogenic response[45–47]. These are a highly conserved group of enzymes, and increased ADAR1 expression and A-to-I editing has been observed in many cancers[48, 49]. In humans, almost all ADAR1 activity occurs within Alu SINEs and Alu editing index (AEI) measures ADAR1-mediated Alu RNA editing[47]. To study the role of IR-Alus and ADAR1 expression in the cancer cells treated with CME inhibitors, we compared the expression levels in our experiments. Interestingly, ADAR1 expression significantly reduced by DNMTi-HDACi in GP5d, OE19 and GP5d p53-KO cells (**Fig. 5e**). We next analyzed AEI to measure the effect of reduced ADAR1 expression on Alu RNA editing. In agreement with reduced ADAR1 expression, we observed significant decrease in AEI in DNMTi-HDACi treated GP5d, OE19 and GP5d p53-KO cells compared to DMSO control (**Fig. 5f**). Moreover, the expression of RIG-I inflammosome, a known cytoplasmic dsRNA sensor[50, 51], significantly increased upon co-inhibition of DNMT and HDAC in all three cell lines (**Fig. 5g**). However, we observed cell type-specific distribution of genomic loci for Alu RNA editing in three cell lines. In GP5d cells, two thirds of the Alu editing-induced mismatch sites were observed within the exonic Alu elements, and the genomic distribution remained similar between DMSO- and DNMTi-HDACi-treated cells (**Fig. 5h**). Intronic Alus represented 29.23% and 35.27% of Alu RNA editing events in DMSO-treated OE19 and GP5d p53-KO cells, and the proportion increased to 41.92 and 44.19% upon DNMT and HDAC co-inhibition, respectively. In contrast, Alu RNA editing in exonic regions decreased from 53.41% to 28.67% and 47.36% to 26.43% by DNMTi-HDACi in OE19 and GP5d p53-KO cells, respectively (**Fig. 5h**). Our results showed that co-inhibition of DNMT and HDAC induces IR-Alu expression along with downregulation of ADAR1. Consequently, we observed decreased Alu RNA editing, suggesting that the DNMTi-HDACi treatment results in increased levels of immunogenic dsRNA.

We further analyzed AEI for all CMEi treatments to study the effect of inhibition of distinct CMEs on Alu RNA editing. Interestingly, HDACi had cell type-specific effect on Alu editing, resulting in increased AEI in both wild-type and p53-KO GP5d cells, whereas in OE19 cells decrease in AEI was observed (**Supplementary Figure 7a**). SETDB1 inhibition, on the other hand, induced a strong increase in Alu editing in all three cell lines (**Supplementary Figure 7a**). Interestingly, GP5d p53-KO cells showed almost two-fold higher Alu RNA editing in intronic regions compared to GP5d cells upon DNMTi-HDACi treatment (**Supplementary Figure 5f**). We also observed cell type-specific changes in the distribution of Alu RNA editing events between GP5d and OE19 cells by distinct CME inhibitor treatments (**Supplementary Figure 7b**). Inhibition of different CMEs results in distinct genomic Alu SINEs being targeted for Alu RNA editing events, and Alu RNA editing in genic regions is associated with retained introns, extended UTRs and making non-reference transcripts from known genes[52]. Collectively, inhibition of CMEs induced changes in Alu RNA editing efficiency with impact on TE-derived transcripts.

### Inhibition of SETDB1 derepresses cell-type specific TEs

SETDB1 amplification is associated with immune escape in human tumors[20]. To study SETDB1i-mediated TE repression in distinct cancer types, we compared TE derepression by SETDB1 inhibition in GP5d, OE19 and GP5d p53-KO cells to publicly available data from SETDB1-KO A375 skin melanoma cells. TEs from LTR subfamilies represented more than half of the TEs derepressed by SETDB1-KO A375 cells[20], and we also observed overrepresentation of LTR subfamilies in differentially expressed TE subfamilies by SETDB1 inhibition (**Fig. 6a, Supplementary Data 4**). Comparative analysis of the expression of TE subfamilies among four different cell lines from melanoma, colon and esophageal cancers revealed cell type-specific derepression of TE subfamilies by SETDB1 KO or inhibition. For example, LTRs from MER52, MER34 and LTR57 families were derepressed in A375 cells, whereas LTR22, LTR73 and MER72B were derepressed in GP5d cells (**Supplementary Figure 8a**). Similarly, LTRs from LTR7, HERV3-int, HERV4-int families and L1ME3E LINEs were derepressed in OE19 cells and LTRs from HERV9 and LTR12 families in GP5d p53-KO cells (**Supplementary Figure 8a**). Generally, the effect of SETDB1 KO/inhibition on differential TE expression was stronger in p53 mutant OE19 and p53-KO GP5d cells compared to wild-type p53-harboring GP5d and A375 cells (**Fig. 6b**). Furthermore, derepressed TEs in different cell types showed distinct distribution to major TE classes. LTRs represented 43% of the TEs derepressed by SetDB1 KO in A375, while SINEs represented 42% of the TEs derepressed by SETDB1 inhibition in GP5d cells. LINEs represented a larger proportion in p53-mutant OE19 and GP5d p53-KO cells (**Fig. 6c**). Around 35% of derepressed TEs in GP5d and GP5d p53-KO cells were Alu SINEs, whereas Alus represented only 14% of the derepressed TE in A375 cells (**Fig. 6d**). Overlap analysis showed that distinct TE loci were derepressed by SETDB1 KO/inhibition in different cell types, but stronger overlap was observed among derepressed TEs in p53 mutant OE19 and GP5d p53-KO cells (**Fig. 6e**). As expected from the small overlap observed between derepressed TEs from different cell types, we also observed cell type-specific TF motif enrichment among derepressed TEs. KRAB-ZFPs such as ZNF460 were enriched in GP5d and A375 cells, while ZNF135 motif was enriched in GP5d cells (**Fig. 6f**). TFs motifs for PRDM9, ZNF384, ZNF257 and MEF2 showed stronger enrichment in OE19 and GP5d p53-KO cells (**Fig. 6f**). Moreover, comparison of the effect of SETDB1 inhibition on AEI showed increased AEI in SETDB1 inhibited cells. Notably, increase in AEI was at least two-fold higher in OE19 cells as compared to GP5d wild-type and p53-KO cells (**Fig. 6g**). Interestingly, intergenic and exonic Alus contributed to higher Alu RNA editing in p53-mutant OE19 and p53-KO GP5d cells, whereas intronic Alus showed a larger proportion of the mismatch sites with RNA editing in GP5d cells (**Fig. 6h**).

**Fig. 6.**
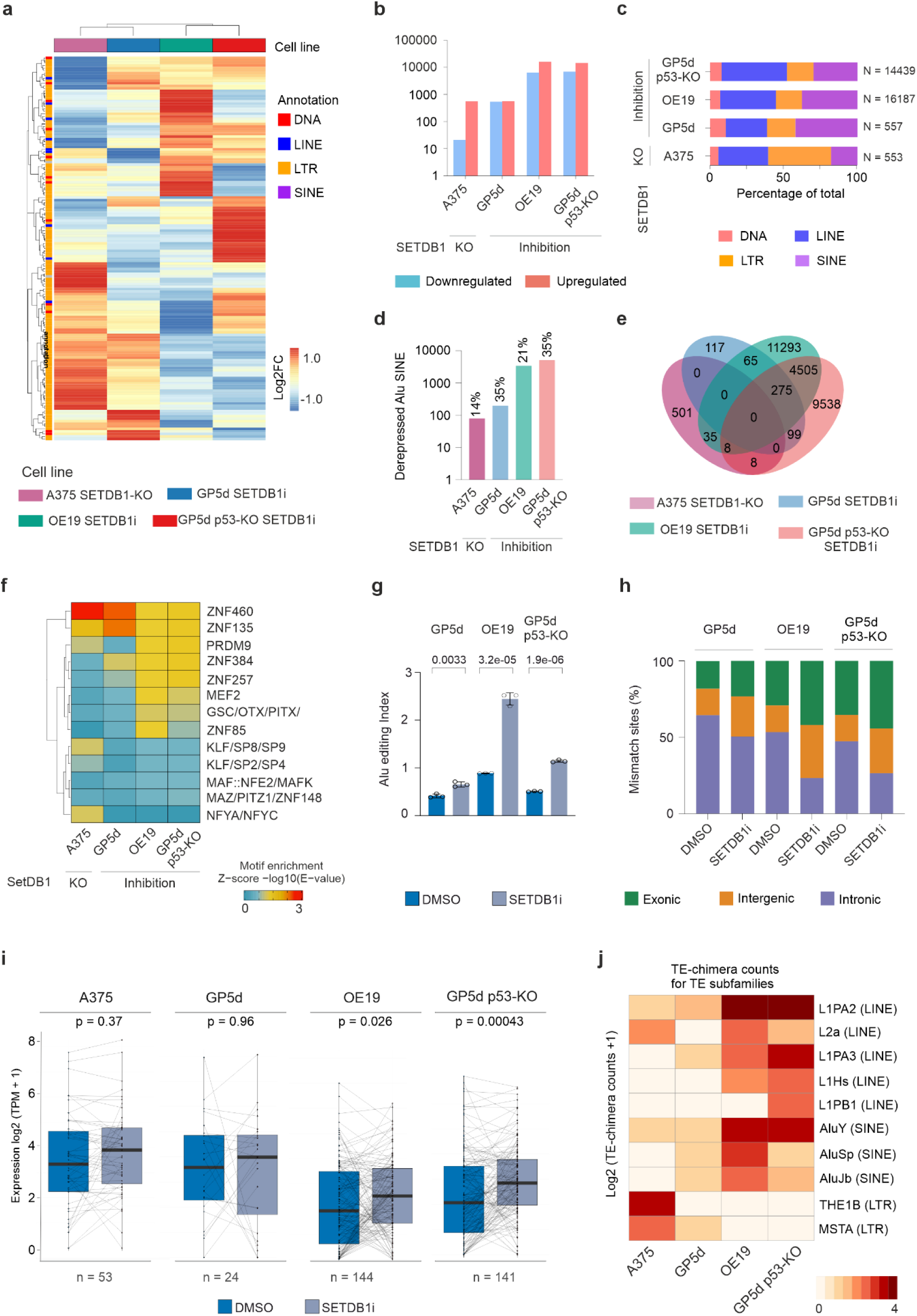
Cell-type specific TEs derepressed by SETDB1 inhibition. a) Comparison of SETDB1 KO/inhibition-induced changes in the expression of TE subfamilies between four cell lines. Differential expression analysis of RNA-seq data was performed by DESeq2. All TE subfamilies with absolute Log2FC > 1.5 (treatment vs DMSO control or A375 KO vs A375) and adjusted p-value < 0.05 in at least one CMEi treatment in one cell line were filtered out and their expression changes in Log2FC was plotted. Rows and columns are clustered with hierarchical clustering. b) Number of differentially expressed TE loci by SETDB1 KO/inhibition in A375, GP5d, OE19 and GP5d p53-KO cells. c) The proportion of TEs derepressed by SETDB1 KO/inhibition belonging to the major TE classes among the derepressed TEs by in A375, GP5d, OE19 and GP5d p53-KO cells. d) Number of Alu SINEs derepressed by SETDB1 KO/inhibition in A375, GP5d, OE19 and GP5d p53-KO cells. Number above the bar represents the percentage of Alu SINEs in total derepressed TEs. e) Overlap of derepressed individual TE loci by SETDB1 KO/inhibition in A375, GP5d, OE19 and GP5d p53-KO cells. f) Motif enrichment analysis for TEs derepressed by SETDB1 KO/inhibition in A375, GP5d, OE19 and GP5d p53-KO cells. Motif enrichment analysis was performed as described in Fig. 3a. g) Bar plots showing Alu Editing Index (AEI) calculated by using RNAeditingIndexer[47] tool for A375, GP5d, OE19 and GP5d p53-KO cells with and without SETDB1 inhibition/KO. h) Distribution of Alu editing sites in A375, GP5d, OE19 and GP5d p53-KO cells with and without SETDB1 inhibition/KO. i) SETDB1 inhibition/KO increases expression of TE-chimeric transcripts. Boxplots showing the expression of TE-chimeric transcripts in A375, GP5d, OE19 and GP5d p53-KO cells with and without SETDB1 inhibition/KO. Expression of TE-chimeric transcripts were determined by TEprof2 analysis pipeline[23]. P values were calculated with a two-sided, Wilcoxon test (n = 53, 24, 144 and 141 differentially expressed TE-chimeric transcripts in A375, GP5d, OE19 and GP5d p53-KO cells, respectively). j) Analysis of TE subfamilies from which the TE-chimeric transcripts are derived from upon SETDB1i inhibition/KO. The counts for TE-chimeric transcripts are log2 transformed.

Earlier studies have shown that TEs epigenetically deregulated in cancer can serve as cryptic promoters and splice into nearby protein coding genes to form TE-chimeric transcripts[21–23]. These transcripts can produce TE-chimeric peptides that can serve as cancer-specific antigens and thus exploited to target cancer cells[22, 23]. To study the effect of SETDB1 inhibition on TE-chimeric expression, we performed TEProf2[23] analysis on RNA-seq data in GP5d, OE19, GP5d p53-KO cells and also from A375 melanoma cells[20]. We observed that SETDB1 inhibition significantly increased the expression of TE-chimeric transcripts compared to DMSO-treatment in OE19 and GP5d p53-KO cells (**Fig. 6i**). The effect was strongly affected by functional p53 status, as fewer TEs contributes to chimeric transcripts formation A375 (n = 53) and GP5d (n = 24) cells with wild-type p53 compared to p53-mutant OE19 (n = 144) and GP5d p53-KO cells (n = 141) (**Fig. 6i**). Interestingly, different TEs contributed to the formation of TE-chimeric transcripts in a cell type-specific manner. For example, THE1B LTRs formed TE-chimeric transcripts in A375 cells and L1PB LINEs in GP5d p53-KO cells (**Fig. 6i**). LINEs from L2a, L1PA3 and L1Hs families and AluJb and AluSp SINEs formed more chimeric transcripts in OE19 and GP5d p53-KO cells (**Fig. 6j**). L1PA2 LINEs and AluY SINEs formed TE-chimeric transcripts in all four cell lines, but the number of these transcripts was much higher in OE19 and GP5d p53-KO cells compared to A375 and GP5d cells (**Fig. 6j**). Collectively, we observed stronger expression of LINE- and SINE-derived TE-chimeric transcripts from SETDB1-inhibited p53-mutant OE19 and p53-KO GP5d cells compared to the cells harboring wild-type p53. As Alu RNA editing in genic regions associated with intron retention and UTR extension[52], we also observed Alu-derived TE-chimeric transcripts in SETDB1 inhibited cells. In agreement with AEI, we observed more Alu-derived TE-chimeric transcripts in OE19 and GP5d p53-KO cells compared to GP5d cells (**Fig. 6j**). Furthermore, gene set enrichment analysis (GSEA) in SETDB1-inhibited GP5d, OE19 and GP5d p53-KO cells revealed enrichment of inflammatory response and interferon-alfa response (**Supplementary Figure 9a**). We also observed increased expression of the interferon stimulated genes[53] by SETDB1 inhibition (**Supplementary Figure 9b**). Collectively, our results demonstrate that SETDB1 KO or inhibition results in cell type-specific depression of TEs, and that the derepressed TEs contribute to TE-chimeric transcript formation to activate immune response in cancer cells.

### Co-inhibition of DNMT and HDAC increases TE-chimeric transcript expression and activates inflammatory response

After observing TE-chimeric transcript expression induced by SETDB1 inhibition, we next analyzed the effect of DNMT and HDAC co-inhibition on TE-chimeric transcripts expression by performing TEProf2[23] analysis on RNA-seq data in colon and esophageal cancer cells. Unlike SETDB1i, DNMTi-HDACi significantly increased the expression of TE-chimeric transcripts in all the three cell lines (**Fig. 7a**), but similar to SETDB1i, the effect was strongly dependent on p53 activity as very few TE-chimeras were expressed by DNMTi-HDACi in GP5d cells (n = 35) compared to p53-mutant OE19 (n = 175) and GP5d p53-KO cells (n = 193) (**Fig. 7a**). Thus, an inverse correlation was observed between functional p53 activity and DNMTi-HDACi induced TE-chimeric transcript formation (**Fig. 7a**). In agreement with earlier reports[21, 22], majority of the TE-chimeric transcripts were derived from LTR12C (**Fig. 7b**). However, we also observed cell type-specificity in TE derived chimeric transcript formation suggesting distinct epigenetic mechanisms. For example, L1PA2, L2a and AluSx subfamilies formed higher number of TE-chimeric transcripts in OE19 cells and AluY SINEs and LTR12D elements in GP5d-null cells (**Fig. 7b**). Genome browser snapshots show TE-chimeric transcripts from LTR12C located near the PARP16 gene for GP5d and p53-KO cells (**Fig. 7c**). Co-inhibition of DNMT and HDAC resulted in stronger derepression of the LTR12C element and higher number of chimeric transcripts derived from LTR12C with exons 2 and 3 of PARP16 in p53-KO cells compared to GP5d cells (**Fig. 7c**). Genome browser snapshot show TE-chimeric transcripts derived from LTR12E located in the first intron of FBP2 gene for OE19 cells (**Fig. 7c**).

**Fig. 7.**
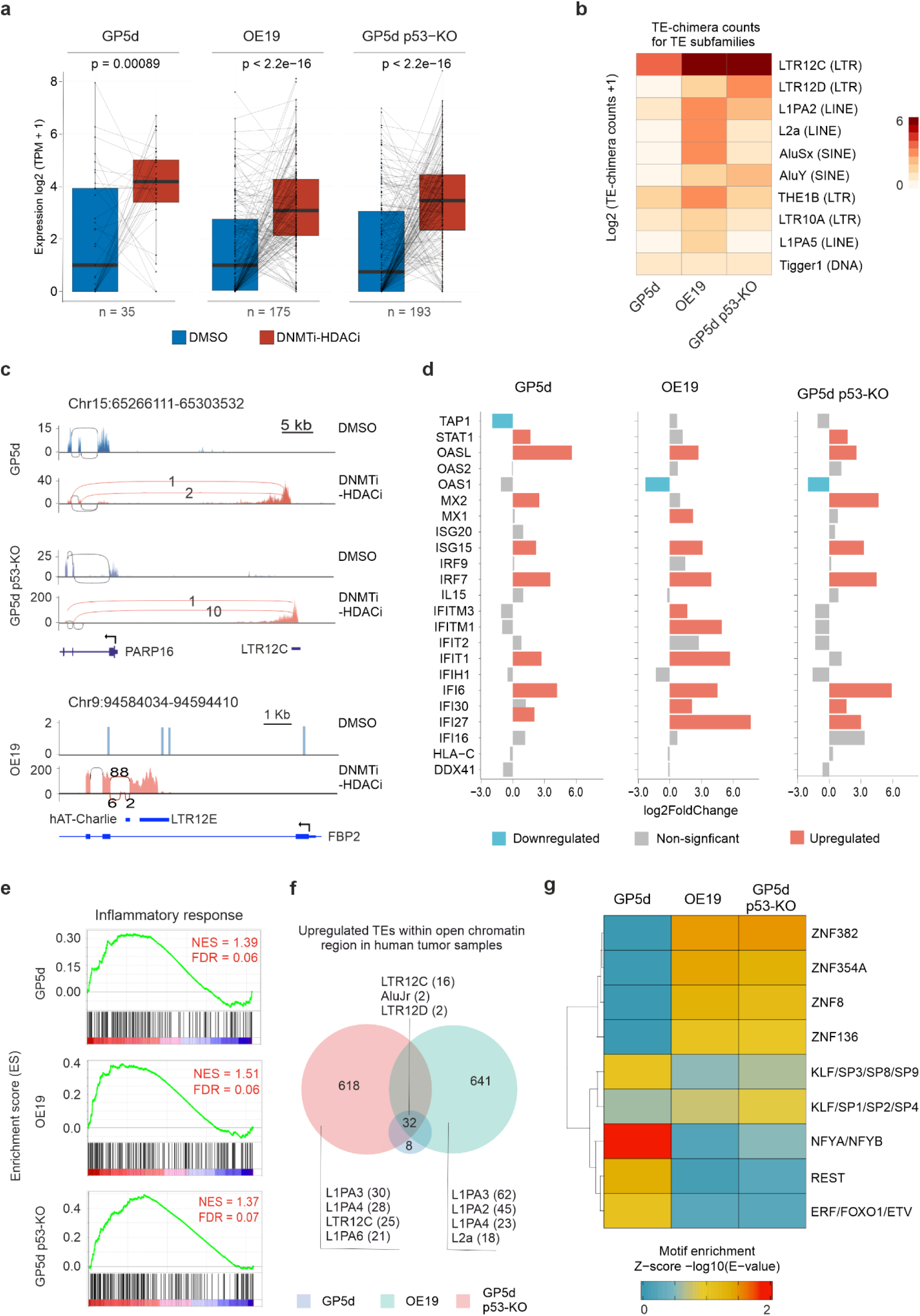
Co-inhibition of DNMT and HDAC increases TE-chimeric transcript expression and activates inflammatory response. a) Co-inhibition of DNMT and HDAC increases expression of TE-chimeric transcripts. Boxplots showing expression of TE-chimeric transcripts in GP5d, OE19 and GP5d p53-KO cells with and without co-inhibition of DNMT and HDAC. P values were calculated with a two-sided, Wilcoxon test (n = 35, 175 and 193 for GP5d, OE19 and GP5d p53-KO cells, respectively). b) Cell type-specific expression of TE subfamilies derived TE-chimeric transcripts by DNMTi-HDACi treatment. The counts for TE-chimeric transcripts are log2 transformed. c) Genome browser snapshot showing LTR12C spliced into second and third exon of PARP16 gene to encode LTR12C-derived TE-chimeric transcript in DNMT and HDAC co-inhibited GP5d (top panel) and GP5d p53-KO cells (middle panel). TE-chimeric transcripts from LTR12C are highlighted in red with average number of TE-chimeric reads from three biological replicates indicated. Strong derepression of LTR12C in GP5d p53-KO cells contributes to higher number of TE-chimeric transcripts. Genome browser snapshot showing LTR12E spliced into second exon of the FBP2 gene to encode LTR12E-derived TE-chimeric transcript from OE19 cells treated with DNMTi-HDACi (lower panel). d) Interferon stimulated genes are upregulated by co-inhibition of DNMT and HDAC. Log2 fold changes in interferon stimulated genes were plotted for DNMTi-HDACi treated GP5d (left panel) OE19 (middle panel) and GP5d p53-KO (right panel) cells. e) GSEA for enrichment of inflammatory response pathway in DNMTi-HDACi-treated GP5d, OE19 and GP5d p53-KO cells. Net enrichment score (NES) and FDR q-value for each pathway is written on top-right corner. f) Overlap analysis of TEs derepressed by DNMTi-HDACi treatment in GP5d, OE19 and GP5d p53-KO cells within open chromatin regions in human tumor samples of the respective cancer types. TCGA ATAC-seq peaks for colon adenocarcinoma (COAD) and esophageal carcinoma (ESCA) were used. Venn diagram represents overlap between derepressed TEs with open chromatin profile in tumor samples. g) Motif enrichment analysis for TEs derepressed by DNMTi-HDACi treatment in GP5d, OE19 and GP5d p53-KO cells and having an open chromatin profile in tumor samples shown in Fig. 7f. Motif enrichment analysis was performed as described in Fig. 3a.

We next compared the differentially expressed genes upon distinct CMEi treatments. In agreement with the changes in TE expression, DNMTi-HDACi treatment had the strongest effect on gene expression in all three cell lines compared to inhibition of the individual enzymes alone (**Supplementary Figure 10a**). Similarly, loss of p53 was associated to stronger changes in gene expression (**Supplementary Figure 10a**). GSEA from the differentially expressed genes induced by the DNMTi-HDACi treatment revealed enrichment of the inflammatory response pathway in both GP5d and GP5d p53-KO cells (**Fig. 7d**). Moreover, the expression of interferon stimulated genes[53] was significantly increased by DNMTi-HDACi in GP5d, OE19 and GP5d p53-KO cells (**Fig. 7e**). Collectively, our GSEA analysis and differential expression of interferon response genes shows that co-inhibition of DNMT and HDAC activates immune response in cancer cells.

Cell type-specificity of the TEs derepressed by co-inhibition of DNMT and HDAC in colon and esophageal cancer cells prompted us to delineate the chromatin state of these derepressed TEs human cancer patient samples. For this, we analyzed the ATAC-seq data for colon adenocarcinoma (COAD) and esophageal carcinoma (ESCA) cancer types using The Cancer Genome Atlas (TCGA) datasets[54]. Importantly, OE19 and GP5d p53-KO specific derepressed L1PA subfamilies were enriched in ESCA and COAD patient samples, respectively (**Fig. 7f**). In addition, the LTR12C and LTR12D elements that were commonly derepressed in all three cell lines also were present within the open chromatin regions of COAD and ESCA patient samples (**Fig. 7f**). Distinct TF motifs were enriched in the derepressed TEs within open chromatin profile, such as motifs from NFY, REST, KLF and SP families in GP5d cells and motifs for zinc finger proteins such as ZNF382, ZNF8, ZNF354A and ZNF136 in OE19 and GP5d p53-KO cells (**Fig. 7g**). Collectively, our results revealed DNMTi-HDACi-induced cell type-specific derepression of TEs, which is associated with cancer type-specific enrichment of TF motifs within the open chromatin regions in TCGA patient tumor samples.

## Discussion

Increasing amount of evidence has shown that epigenetic therapy alone or as an adjuvant to immunotherapy provides better therapeutic output in cancer patients[17, 18, 25]. TE derepression induced by epigenetic therapy plays a key role in making cancer cells sensitive to immunotherapy[18, 22]. However, the epigenetic regulation of the TEs by specific CMEs and in context of p53 which is mutated in majority of human cancers is less well understood. Here, we studied the role of distinct CMEs and p53 in TE regulation in two endodermal origin cancers. We identified TE subfamilies that are under distinct epigenetic regulation and showed that they are transcriptionally activated by inhibition of distinct CMEs in a cell-type specific manner. We found that loss of functional p53 activity is associated with stronger derepression of TEs by different CME inhibition treatments. Moreover, we report that KAP1 binding along with gain of active histone marks is associated with derepression of specific TE subfamilies upon co-inhibition of DNMT and HDAC. Finally, we show that TEs derepressed by inhibition of CMEs contribute to activation of immune response by inducing the expression of IR-Alu SINEs and TE-derived chimeric transcripts.

Our subfamily level analysis of derepressed TEs in three different cell lines revealed that majority of derepressed TE subfamilies are from the LTR class. Previously, it has been reported that although LTRs represents only 8% of the human genome, they account for 39% of TF binding sites in the human genome[55]. Overrepresentation of LTRs in derepressed TE subfamilies is also consistent with our earlier report showing stronger enrichment of LTRs in STARR-seq enhancers in colon and liver cancer cells[5]. TEs are epigenetically silenced by various mechanism such as cytosine methylation of DNA[4, 7], enrichment of repressive histone modifications on nucleosome[4, 8] and recruitment of transcriptional repressors such as KRAB-ZNFs[9, 56] and p53[4, 11]. As TEs have a broad diversity in their GC-content, sequence composition[57] and genomic location, they are bound to be targeted by distinct epigenetic silencing mechanisms. In agreement with this, we observed that (i) distinct TE subfamilies are epigenetically derepressed by inhibition of the specific CMEs in the same cell line, (ii) there is a minimal overlap among derepressed individual TE loci induced upon different CME inhibition treatments in the same cell line, and (iii) TEs derepressed by different CME inhibition treatments show distinct genomic distributions. Of note, we did not observe stronger derepression of TEs by DNMTi compared to HDACi that was previously reported for ovarian cancer cell lines[37], suggesting cancer cell type-specific response. SETDB1 is recruited to TEs by KRAB-ZFP/KAP1 repressive complex[9] or core members of HUSH complex[58, 59] to deposit repressive H3K9me3 marks. Notably, SETDB1i had stronger effects on TE derepression than HDACi or DNMTi alone in all three cell lines, whereas co-inhibition of DNMT and HDAC showed synergistic effect on TE derepression, in agreement with earlier reports[21, 22].

Inhibition of different CMEs in GP5d, OE19 and GP5d p53-KO cells results in derepression of both cell type-specific as well as common TE subfamilies. For example, DNMTi-HDACi induced derepression of LTRs from LTR12 and HERV9 subfamilies in both colon and esophageal cancer cells, whereas LTR10B was specifically derepressed by DNMTi-HDACi in GP5d cells, further supporting our earlier finding of enrichment of LTR10B elements in GP5d STARR-seq enhancers[5]. In general, we did not observe strong representation of interspersed repeats in differentially expressed TE subfamilies, but some of them showed cell type-specific derepression. For example, L1ME3E elements were derepressed in GP5d cells and AluYh9 in OE19 cells by SETDB1 inhibition.

In addition to derepression of TEs upon CME inhibition, we also observed repression/downregulation of TEs and genes by inhibition of CMEs in GP5d, OE19 and GP5d p53-KO cells. This effect was stronger for SETDB1 inhibition compared to inhibition of other CMEs (see Fig. 2A, 2D, S10A). Recently, SETDB1 was reported to control chromatin organization through regulating CTCF binding independent of DNA methylation in mouse embryonic stem cells (mESC)[60]. Loss of SETDB1 increased chromatin accessibility and CTCF binding at rodent-specific B2 SINEs in mESC. Loss of SETDB1 perturbed local chromatin interactions which was also associated with repression and derepression of TEs and genes[60]. Further studies are required to understand chromatin organization changes induced by inhibition of CMEs for better insights into transcriptional regulation of TEs and genes by distinct CMEs.

In the context of TE regulation, p53 has been described as transcriptional repressor as well as transcriptional activator[39, 61], and our results support both of these functions. We observed reduced RNA and protein levels of p53 itself in DNMTi-HDACi-treated GP5d and p53-mutant OE19 cells. In support of the repressive role of p53 in TE regulation, we observed greater number of TE loci derepressed by CME inhibitor treatments in p53-mutant OE19 and p53-KO GP5d cells compared to GP5d cells with wild-type p53. We observed higher percentage of LINEs in TEs loci derepressed by distinct treatments in OE19 and p53-KO GP5d cells compared to GP5d cells, which also supports earlier reported higher expression of LINE-1 in p53-mutant tumors[38]. However, p53 has also been reported to bind closed chromatin[30, 62] to transactivate nearby genes by binding to p53RE containing LTRs[39]. In support of p53 as an activator, we found downregulation of the earlier reported p53-target genes that are regulated by p53-bound LTR10 elements in GP5d cells (**Supplementary Figure 4a-b**). Collectively, our results highlight the context-specificity of p53 function with both repressive and activating roles in TE regulation.

The role of KAP1 in transcriptional regulation is not completely understood[14, 63]. The repressive function of KAP1 is extensively characterized[9, 12, 56], but KAP1 is also recruited to actively transcribed genes and involved in releasing the paused Polymerase II[14, 16]. KAP1 has been shown to interact with H4K16ac-modified histone H4 at the gene promoters[15]. Here, we show KAP1 binding at H4K16ac-enriched LTR12C elements in GP5d and OE19 cells that is also associated with KAS-seq signal for ssDNA in OE19 cells. Thus, association of KAP1 with H4K16ac marks is not only restricted to actively transcribing gene promoters, but occurs at derepressed TE loci as well. LTRs and LINEs enriched with H4K16ac marks have been shown to have enhancer activity in human embryonic stem cells[40]. However, we did not observe such cis-regulatory activity from H4K16ac-marked LTR12C elements in differentiated colon and esophageal cancer cells, suggesting developmentally restricted role for H4K16ac in TE regulation. Interestingly, we observed different epigenetic states for derepressed LTR12C elements in GP5d and OE19 cells, namely that active H3K4me3 mark was increased in the LTR12C elements in DNMTi-HDACi-treated GP5d cells but the levels remained constant in OE19 cells. However, the gain of H3K27ac marks was the major determinant of the transcriptional activation of TEs in both GP5d and OE19 cells. Notably, derepressed LTR12C elements were associated to the transactivation of nearby genes. Collectively, we show that TEs from same subfamily are under cell type-specific epigenetic regulation and reactivated by enrichment of H3K27ac marks by DNMTi-HDACi treatment.

Our expression analysis from GP5d and GP5d KAP1^+/-^ cells upon DNMT and HDAC co-inhibition revealed the dual role of KAP1, functioning as both repressor and activator in TE regulation. In agreement with its known role as TE repressor, loss of one KAP1 allele resulted in derepression of larger number of TE loci and stronger derepression at majority of derepressed TE loci compared to GP5d cells. However, TEs from specific subfamilies such as LTR12C showed weaker derepression in GP5d KAP1^+/-^ cells compared to GP5d cells, which suggests transcriptional activator role for KAP1 in DNMTi-HDACi-treated cells. Furthermore, we observed recruitment of KAP1 at derepressed LTR12C elements in DNMTi-HDACi treated GP5d cells. Thus, our data suggest that chromatin-bound interactome of KAP1 determines its effect on transcription. Further studies are required to understand how the dynamics of KAP1 interactome change by inhibition of distinct CMEs.

TE-derived dsRNA activates immunogenic response[24, 25] and the IR-Alus are the major source of the dsRNA[24, 47]. ADAR1 enzymes edit dsRNA from IR-Alus to inhibit activation immunogenic response[45–47]. Here, we report derepression of the IR-Alus by DNMTi-HDACi treatment with OE19 cells showing greater number of IR-Alus compared to GP5d. Derepressed IR-Alu also gained H3K27ac and KAP1 ChIP-signal in both GP5d and OE19 cells. We also observed downregulation of ADAR1 expression with concomitant upregulation of RIG-I. However, the expression of MDA5 (IFITH1), earlier reported protein sensor for dsRNA[64], was not upregulated by DNMTi-HDACi (**Fig. 7e**). Our comparative analysis revealed that distinct CME inhibitor treatments altered Alu RNA editing efficiency. Interestingly, SETDB1 inhibition resulted in increased AEI along with derepression of Alu SINEs in all three cell lines, but the genomic distribution of Alu RNA editing sites was cell type-specific. Alu RNA editing in genic regions can contribute to intron retention and UTR extension[52]. However, to what extent the changes in Alu RNA-editing efficiencies elicited by the inhibition of CMEs alter protein coding transcriptomes warrants further studies. Collectively, different epigenetic therapies induced changes in Alu RNA editing efficiencies, emphasizing their potential for regulating immunogenic response for improved cancer therapeutics.

Chimeric TE-transcripts derived from derepressed TEs can serve as novel antigens specific for cancer cells. Here, we report that loss of p53 activity was associated with greater number of TE-derived chimeric transcripts induced by SETDB1i and DNMTi-HDACi. In OE19 and GP5d p53-KO cells, SETDB1 inhibition induced chimeric transcripts derived primarily from LINEs and SINEs, whereas LTR12C-derived transcripts were overrepresented in the same cell lines by co-inhibition of DNMT and HDAC. These results suggest that different TE classes are derepressed by inhibition of different CMEs, allowing TE-embedded cryptic promoters to form TE-derived chimeric transcripts. Overrepresentation of LTR12C-derived chimeric transcripts upon DNMTi-HDACi treatment reported here is in agreement with earlier studies[21, 22]. However, we also observed TE-chimeric transcripts from other TE subfamilies induced in a cancer cell type-specific manner. For example, greater number of chimeric transcripts were formed from L1PA2, L1PA3 and L1Hs LINEs in SETDB1i- and DNMTi-HDACi-treated OE19 cells compared to GP5d cells. Such LINE1-derived TE-chimeric transcripts are also supported by earlier evidence of LINE1 anti-sense promoter derived chimeric transcripts in healthy brain[65] and cancer cells[66]. TE-derived chimeric transcripts encode for immunogenic peptides that can be exploited to target cancer cells using immunotherapy[22, 23]. Importantly, our results show that epigenetic therapy-induced TE-chimeric transcripts are cell type-specific and that their expression is higher in p53-mutant cancers compared to p53 wild-type cells. Furthermore, the activation of immune response and upregulation of the interferon stimulated genes upon SETDB1i and DNMTi-HDACi treatment suggest an appealing therapeutic strategy for making the cancer cells more vulnerable and responsive to immunotherapy.

In conclusion, we studied the extent of TE regulation by distinct CMEs and p53 in colon and esophageal cancer. We found that TEs are under distinct epigenetic regulation and that they are released from repression by inhibition of CMEs in a cell type-specific manner. Loss of p53 was associated with stronger derepression of TEs, whereas KAP1 had also TE activating function in DNMT and HDAC co-inhibited colon and esophageal cancer cells. Derepressed TEs gain epigenetic signatures of active enhancers and transactivates nearby genes. TE derepression results in activation of immune response. Thus, a deeper understanding of epigenetic regulation of TEs could yield epigenetic therapies that selectively enhance anti-tumor immunity, paving the way for precision cancer medicine.

## Methods

### Data acquisition

All sequencing data and download links for annotation files used in this study are listed in **Supplementary Data 5** including the relevant references and GEO/ENCODE accessions. A gene annotation GTF file was downloaded from Gencode Release 36 for the reference chromosomes. The GTF file was converted into a BED file with gtfToBed.sh and TSS and gene body BED files were created with a script adapted from ref. [67].

A repeatMasker.txt (2021-09-03) file was downloaded from the UCSC table browser. Only transposable element-derived repeat classes (LINE, SINE, LTR, and DNA) were retained and a file in BED format was created from the table, totaling 4745258 annotated repeats[68]. LTR12C consensus sequence was downloaded from RepBase[69].

GRCh38 chromosome sizes file was downloaded from UCSC.

GRCh38 blacklist BED file ("ENCFF356LFX [https://www.encodeproject.org/files/ENCFF356LFX/]”, release 2020-05-05) was acquired from the ENCODE project.

A genome index was created with bowtie2-build, with chr1-22, X, Y and M fasta files. Alternative, unlocalized and unplaced alternative loci scaffolds were discarded in indexing. Transcription factor motifs were acquired from JASPAR 2022 CORE non-redundant vertebrate annotations[70]. The position weight matrices in MEME format were used for motif enrichment analyses.

TCGA cancer-type specific ATAC-seq peaks for colon adenocarcinoma (COAD) and esophageal carcinoma (ESCA) were acquired from ref [54].

### Cell culture

GP5d (Sigma, 95090715) and GP5d p53-KO cells were cultured in DMEM (Gibco, 11960-085) supplemented with 10% FBS (Gibco, 10270106), 2 mM L-glutamine (Gibco, 25030024) and 1% penicillin-streptomycin (Gibco, 15140122). OE19 (Sigma, 96071721) cells were cultured in RPMI (Gibco, 31870) supplemented with 10% FBS (Gibco, 10270106), 2 mM L-glutamine (Gibco, 25030024) and 1% penicillin-streptomycin (Gibco, 15140122).

All cell lines were directly obtained from Sigma and low-passage cells were used in all experiments. All cell lines tested negative for mycoplasma contamination upon purchase and were routinely checked as per standard good laboratory practice.

### Generation of GP5d KAP1 ^+/-^ cell line by genome editing

The KAP1^+/-^ GP5d cell line was generated by CRISPR-Cas9 targeting of exon 2 of the KAP1 gene using Alt-R CRISPR-Cas9 from Integrated DNA Technologies. Briefly, Equimolar ratios of target-specific crRNA (**Supplementary Data 6**) and ATTO550-tracrRNA (IDT, 1075928) were annealed and RNP complex were constituted from Alt-R CRISPR-Cas9 (IDT, 1081060; 1000 ng per 200,000 cells) and target-specific sgRNA (250 ng per 200,000 cells). RNP complex transfected to early passage GP5d cells by using CRISPRMAX (Life Technologies, CMAX000003) according to manufacturer’s protocol. The next day, atto550+ cells were FACS sorted, and single-cell colonies were cultured to produce a clonal KAP1^+/-^ cell line. The clonal cells lines were screened for KAP1 deletion and clones were verified by Sanger sequencing using primers flanking deletion site (**Supplementary Data 6**).

### ChIP-seq

ChIP-seq was performed as previously described[30] by using the following antibodies: KAP1 (Bethyl Laboratories, A300-274A), H3K27ac (Diagenode, C15410196), H3K4me3 (Diagenode, C15410003), H4K16ac (Diagenode, C15200219). Each ChIP-seq reaction was performed by using 2 μg of antibody. In brief, GP5d and OE19 cells were cross-linked for 10 minutes at room temperature by using formaldehyde (Sigma, F8775). Sonicated chromatin was centrifuged, and the supernatant was used to immunoprecipitated DNA using Dynal-bead coupled antibodies. Immunoprecipitated DNA was purified and used for ChIP-seq library for Illumina sequencing. ChIP-seq libraries were single-read sequenced on NovaSeq 6000.

### CMEs inhibitor treatment and RNA-seq

DNMT inhibition in GP5d, OE19 and GP5d p53-KO cells were performed as described earlier[21]. Cells were seeded in 6 well plate and treated with 500 nM/L DAC (MedChemExpress, HY-A0004). Media containing DAC containing were replaced each day for three days. Cells were harvested after 72 hours for RNA isolation.

HDAC inhibition in the three cell lines was performed as described earlier[21]. Cells were treated with 500 nM/L SB939 (MedChemExpress, HY-13322) for 18 hours and cells were harvested for RNA isolation.

SETDB1 inhibition in the three cell lines was performed as previously described[71]. Cells were treated with 500 nM/L Mitramycin A (MedChemExpress, HY-A0122) for 24 hours and cells were harvested for RNA isolation.

Co-inhibition of DNMT and HDAC in OE19 and GP5d p53-KO cells were performed as described in ref[5, 21]. Cells were treated with 500 nM/L DAC. Media containing DAC containing were replaced each day for three days. Cells were treated with 500 nM/L SB939 for 18 hours and collected for RNA isolation and ChIP-seq. For DMSO control, cells were treated with DMSO (Fisher, BP231). Media containing DMSO were replaced each day for four days. Cells were collected for RNA-isolation after 4 days.

RNeasy Mini kit (Qiagen) was used to isolate total RNA from different CMEs or DMSO treated cells. RNA-seq libraries were prepared using 500 ng of total RNA by using KAPA stranded RNA-seq kit for Illumina (Roche) as per manufacturer’s instruction. All RNA-seq samples were sequenced paired-end on NovaSeq 6000 (Illumina).

### ChIP-seq analysis

The ChIP-seq reads were mapped with bowtie2 v.2.4.1 (bowtie2 --very-sensitive) to the reference human genome (hg38/GRCh38)[72]. Duplicates were removed by using Picard v.2.23.4 (MarkDuplicates -REMOVE_DUPLICATES false -ASSUME_SORT_ORDER coordinate)[73]. Samtools v.1.7 was used to filter reads with MAPQ smaller than 20 and remove marked duplicates (samtools view -F 1024 -b -q 20)[74]. Peaks were called with MACS2 v.2.2.7.1[75]. Peaks overlapping with ENCODE blacklisted regions were removed from the Peak and summit files with bedtools v.2.29.2 (bedtools subtract -A). RPKM-normalized bigwig file was prepared by using deepTools v.3.5.0 (bamCoverage —binSize 50 --normalizeUsing RPKM)[76]. Pearson correlation analysis between the biological replicates for ChIP-seq is shown in **Supplementary Figure 11a**.

### ATAC-seq analysis

GP5d ATAC-seq data were acquired from ref. [5] and OE19 ATAC-seq data wer acquired from ref. [43]. The ATAC-seq reads were mapped with bowtie2 v.2.4.1[72] (--very-sensitive) to the reference human genome (hg38/GRCh38). Reads mapped to the mitochondrial genome were removed with removeChrom.py script[77] (https://github.com/jsh58/harvard/blob/master/removeChrom.py). Duplicate reads were removed by using Picard v.2.23.4 (MarkDuplicates -REMOVE_DUPLICATES false - ASSUME_SORT_ORDER coordinate)[73] and insert sizes were analyzed by using CollectInsertSizeMetrics. Samtools v.1.7 was used to remove marked duplicates and filter reads with MAPQ smaller than 10 (samtools view -F 1024 -b -q 10). ATAC-seq peaks were called with MACS2 v.2.2.7.1 (macs2 callpeak -f BAMPE -g hs—keep-dup all). Removal of peaks overlapping blacklisted region and preparation of a RPKM normalized bigwig was performed as in ChIP-seq data processing.

### CUT&TAG analysis

OE19 CUT&TAG data for H3K27me3, H3K4me1, RNA-Pol II and Serine-5-phospohorylated RNA-Pol II was acquired from ref. [43]. The CUT&TAG paired-end reads were mapped with bowtie2 v.2.4.1[72] to the reference human genome by using following parameters: --very-sensitive --no-mixed --no-discordant -I 10. Duplicate reads were removed by using Picard v.2.23.4 (MarkDuplicates -REMOVE_DUPLICATES false - ASSUME_SORT_ORDER coordinate)[73]. Samtools v.1.7 was used to filter reads with MAPQ smaller than 20 and remove duplicates (samtools view -F 1024 -b -q 20)[74]. RPKM-normalized bigwig file was prepared by using deepTools v.3.5.0 (bamCoverage --binSize 50 --normalizeUsing RPKM)[76]. RPKM normalized bigwig files were used to plot CUT&TAG signal.

### KAS-seq analysis

OE19 KAS-seq raw data were downloaded under ENA accession “PRJEB50427” (“ERR8135308”). KAS-seq reads were aligned to the hg38 genome assembly using bowtie2 v.2.4.1 (--very-sensitive). Duplicate reads were removed by using Picard v.2.23.4 (MarkDuplicates -REMOVE_DUPLICATES false - ASSUME_SORT_ORDER coordinate)[73]. Samtools v.1.7 was used to remove marked duplicates and filter reads with MAPQ smaller than 20 (samtools view -F 1024 -b -q 20). Preparation of a RPKM normalized bigwig was performed as in ChIP-seq data processing.

### TEtranscripts and Telescope RNA-seq analysis

The RNA-seq reads were mapper with STAR v.2.5.3a by using the SQuIRE pipeline v.0.9.9.92[32]. SQuIRE alignment output was used to TE subfamilies expression measurement by using TEtranscripts v.2.2.1[33] with following flags: –mode multi –stranded reverse. DESeq2 v.1.32.0[78] was used for differential expression analysis of TE subfamilies. Telescope v.1.0.3.1[34] analysis was performed on SQuIRE alignment output by using the “telescope assign” command. DESeq2 v.1.32.0 was used for differential expression analysis for individual TE loci. GREAT v.4.0.4[79] was used to assign nearby gene for TE loci.

To compare TE subfamily expression changes between cell lines (**shown in Fig. 1c and 6a**), TE subfamilies with strong transcriptional changes were filtered out. TE subfamilies with absolute Log2FC of more than 2.5 (treatment vs DMSO control) and adjusted p-value less than 0.05 in at least one CMEs treatment were filtered out and expression changes in Log2FC was plotted by using Pheatmap[80]. Rows and columns are clustered with hierarchical clustering. Heatmap to compare TE subfamilies expression changes for SETDB1 inhibited/KO cell lines (**shown in Fig. 6a**) was plotted as described in **Fig. 1c** by filtering out TE subfamilies with absolute Log2FC of more than 1.5 (treatment vs DMSO control) and adjusted p-value less than 0.05.

### IR-Alu expression analysis

The IR-Alu for human genome reported in ref. [24] was shared by Dr. Parinaz Mehdipour and genomic coordinates were converted from hg19 to hg38 assembly by using liftOver tool[81]. DESeq2 normalized Telescope expression counts were extracted for all IR-Alus and actively transcribed IR-Alu were filtered out (Total sum of RNA-seq reads for DMSO and DNMT-HDACi ≥5) and their expression were compared for GP5d and OE19 cells with and without DNMTi- HDACi treatments.

### Alu editing index analysis

The RNA-seq reads were mapped to human reference genome with STAR v.2.7.5a with parameter recommended in ref. [47] (--outFilterMatchNminOverLread 0.95). Duplicate reads were removed by using Picard v.2.23.4 (MarkDuplicates -REMOVE_DUPLICATES true - ASSUME_SORT_ORDER coordinate). Deduplicated bam files were sorted by using Samtools v.1.7. Alu editing index analysis was performed by using RNAeditingIndexer[47]. A to G Editing counts for exons, intron and intergenic regions were used to calculate the distribution of Alu editing sites.

### Chimeric transcript analysis

TE-chimeric transcripts analysis was performed by using TEprof2 pipeline[23]. The RNA-seq reads were mapped to human reference genome with STAR v.2.7.5a and then assemble files by using stingtie v.1.3.3. TEprof2 v.0.1 was used to identify transcript overlapping with TEs and transcripts from GENCODE v.25. TE-chimeric transcripts were identified using TEProf2 pipeline with default parameter. TE-chimeric transcripts expressed at least two replicates either in control or in treatment samples were selected for downstream analysis. Mean transcript expression was compared between DMSO control and DNMTi-HDACi cells for three cell lines.

### Western blotting

Cells were lysed in RIPA buffer containing 1mM DTT (Thermo scientific, 20290) and protease inhibitor (Roche, 11873580001). Protein samples (50 ug for each sample) were denatured by using 6x SDS-Laemmli buffer (Fisher, 50-103-6570) at 95–100°C for 5 min. Proteins were separated by SDS-PAGE using acrylamide gels (Biorad, 4569033) and transferred to PVDF membrane (Thermo scientific, 88518). Membrane was blocked in in 5% skimmed milk containing 1x TBST buffer and incubated with the following primary antibody: p53 (Santa Cruze Biotechnology, sc-126, 1:1000), KAP1 (Bethyl Laboratories, A300-274A, 1:1500) and GPADH (Santa Cruze Biotechnology, SC-47724, 1:1000). As secondary antibody, goat anti-mouse IgG (Bio-Rad, 5178-2504, 1:5000) and mouse rIgG (Bio-RAD, 5196-2504,1:5000) was used. PVDF Membranes were imaged using Image Studio Lite (Odyssey CLx imager, Li-COR Biosciences). Uncropped images for all western blots shown in **Supplementary Figure 12a**.

### RNA-seq analysis

FASTQC was used for the quality control of raw sequencing FASTQ files[82] (http://www.bioinformatics.babraham.ac.uk/projects/fastqc/). RNA-seq reads were aligned to the human reference genome using STAR aligner v.2.7.5c with default parameters[83] and Samtools v.1.7 was used to sort bam files. Gene counts were quantified by using HTSeq-count v.0.11.2[84]. DESeq2 v.1.32.0 was used to identify differential expressed genes for each CMEs treatment as compared to DMSO control, a threshold of |log2FoldChange| > 1.5 and adjusted p-adj < 0.05 was applied. Gene Set Enrichment Analysis was performed by using GSEA v.4.1.0[85].

### TCGA ATAC-seq analysis

Cancer type-specific ATAC-seq peak sets for COAD and ESCA TCGA cancer types were acquired from ref. [86]. The overlap analysis with derepressed TEs was performed by using bedtools v.2.29.2.

### KAS-seq, ChIP-seq read alignment to LTR12C consensus sequence

KAP1 ChIP-seq reads from DNMT and HDAC co-inhibited GP5d cells were first aligned to the reference genome. Unique reads mapped to 499 derepressed LTR12C elements were extracted and mapped to LTR12C consensus sequence to obtain compiled reads. Bwtools v.1.0 (bwtools aggregate) was used to plot compiled KAP1 ChIP-seq signal at LTR consensus sequence[87].

For OE19, KAS-seq and KAP1 ChIP-seq were first aligned to the reference genome. Unique reads mapped to 861 derepressed LTR12C elements were extracted and mapped to LTR12C consensus sequence to obtain compiled reads. Bwtools v.1.0 (bwtools aggregate) was used to plot compiled signal for KAS-seq, KAP1 ChIP-seq at LTR12C consensus sequence.

### Motif analyses

Motif enrichment analysis at derepressed TEs was performed using AME from MEME suite v. 5.0.2 with shuffled sequences as background (ame --control --shuffle--)[88]. JASPAR 2022 CORE non-redundant vertebrate motif annotations were used as input for the motif file. Motif clustering data were acquired from Viestra et al.[44]. The E-values were –log10 transformed by using R and for each treatment, minimal E-value for individual motifs were selected and the columns were scaled using the R scale function (center = F) and plotted.

### Statistical analysis and plots

All statistical analyses were performed by using R v.4.1.2[89] and GraphPad prism v.9. Genomic annotation for derepressed TEs was performed by using ChIPseeker[90]. Boxplots were prepared by using ggplot2 v.3.3.6 from the Tidyverse suite v.1.3.1[91]. Heatmap for the motif enrichments were plotted with ComplexHeatmap v.2.8.0[92]. Heatmap in **Fig. 1c** and **Fig. 6a** was plotted by using Pheatmap[80]. Correlation analysis between replicates was performed by using multiBigwigSummary v.3.1.3 and heatmaps were plotted using plotCorrelation v.3.1.3[93]. Illustrations were created with BioRender.com. Heatmaps and average profile plots were plotted by using deepTools[93]. Genome browser snapshots for TE-chimeric transcripts were plotted by using ggshashmi v.1.1.5[94]. All motif enrichment heatmaps and genomic annotation bar graphs were plotted by using previous deposited R scripts from ref. [5].

## Data availability

Data generated in this study has been deposited in the GEO database under accession “GSE254242”.

The publicly available data was accessed as follows: GP5d ChIP-seq data for H3K27ac (“GSM5454417”), H3K27me3 (“GSM5454428”), and p53 (“GSM5454412”) were acquired from GEO database under accession “GSE180158”. GP5d ATAC-seq (“GSE221051”), ChIP-seq for H3K4me3 ("GSM6841187”), rabbit IgG ("GSM6841190”) and mouse IgG ("GSM6841189”), and RNA-seq for DMSO treated (“GSM6841203”, “GSM6841204”, “GSM6841205”) and DNMTi-HDACi treated GP5d cells (“GSM6841206”, “GSM6841207”, “GSM6841208”) was acquired from GEO database under accession “GSE221053”. GP5d H3K4me1 (“GSM1240814”) was obtained with the GEO accession “GSE51234”. OE19 ATAC-seq (“ERR1698333”) was downloaded with accession code ERX1767841. OE19 KAS-seq (“ERR8135308”) was acquired from accession code “E-MTAB-11356”. OE19 CUT&TAG data for H3K27me3 (ERR8105268), H3K4me1 (ERR8105270), RNA PolII (ERR8105275) and RNA PolIISp5 (ERR8105277) was obtained from ENA accession “E-MTAB-11356”. A375 RNA-seq (“GSM5320279”, “GSM5320280”, “GSM5320281”) and A375 SETDB1-KO RNA-seq (“GSM5320275”, “GSM5320276”, “GSM5320277”) was acquired from GEO accession “GSE155972”.

## Code availability

No specific code used in the data analysis.

## Acknowledgements

We thank HiLIFE research infrastructures including the FIMM NGS Genomics laboratory at the University of Helsinki. We thank the Center for Scientific Computing (CSC), Finland, for the computational infrastructure, Professor Lauri Aaltonen’s laboratory facilities for genomics work. BS was supported by the Academy of Finland (317807, 320114, 346065), Finnish Cancer Foundation, Sigrid Jusélius Foundation, Jane and Aatos Erkko Foundation and iCAN Digital Precision Cancer Medicine Flagship (320185). We thank Dr. Parinaz Mehdipour for sharing IR-Alu sequences for human genome. We thank Dr. Liangru Fei and Dr. Jihan Xia for technical assistance with NGS library preparation. We thank Konsta Karttunen and Nikita Poddar for technical help in software installation and Pinja Perkkiö for laboratory assistance.

## Author Contributions

DP and BS conceived the study. BS supervised the study. DP performed experiments and genomics data analysis. VT helped with the gene expression analysis. PP helped with interpretation and presentation of the data. DP and BS wrote the manuscript with contributions from all authors.

## Competing interests

The authors declare no competing interests.

## Notes

### Competing Interest Statement

The authors have declared no competing interest.

### Summary of Updates

Manuscript discussion section updated; Figure 6 revised.

